# Genome-wide analysis of DNA-PK-bound MRN cleavage products supports a sequential model of DSB repair pathway choice

**DOI:** 10.1101/2022.12.07.519501

**Authors:** Rajashree A. Deshpande, Alberto Marin-Gonzalez, Taekjip Ha, Tanya T. Paull

## Abstract

The Mre11-Rad50-Nbs1 (MRN) complex recognizes and processes DNA double-strand breaks for homologous recombination by performing short-range removal of 5ʹ strands. Endonucleolytic processing by MRN requires a stably bound protein at the break site—a role we postulate is played by DNA-dependent protein kinase (DNA-PK) in mammals. Here we interrogate the sites of MRN-dependent processing by isolating and sequencing DNA-PK-bound DNA fragments that are products of MRN cleavage. These intermediates are generated with highest efficiency when DNA-PK is catalytically blocked, yielding products within 200 bp of the break site, whereas DNA-PK products in the absence of kinase inhibition show much greater dispersal. Use of light-activated Cas9 to induce breaks facilitates temporal resolution of DNA-PK and Mre11 binding, showing that Mre11 and DNA-PK both bind to DNA ends before release of DNA-PK-bound products. These results support a sequential model of double-strand break repair involving collaborative interactions between homologous and non-homologous repair complexes.

## Introduction

Double-strand breaks (DSBs) in genomic DNA are sensed by several evolutionarily conserved protein complexes that protect and repair the lesions. The most abundant of these in mammalian cells is DNA-dependent protein kinase (DNA-PK) composed of the catalytic kinase subunit (DNA-PKcs) and the DNA end-binding heterodimer Ku. The Ku component of DNA-PK appears in all kingdoms of life while DNA-PKcs is primarily found in higher eukaryotes^1^. Together, these factors protect DNA ends from nonspecific nuclease degradation and facilitate the recruitment of other factors involved in non-homologous end joining (NHEJ)^2^.

NHEJ is considered to be the major pathway for double-strand break repair in mammals, although replication-dependent repair pathways that rely on homologous recombination for the resolution of processing intermediates are also essential for cell viability in dividing cells^3,4^. Homology-driven pathways require that DSB ends first be resected to remove hundreds of nucleotides from the 5ʹ strand, in order to facilitate loading of the Rad51 recombinase on the 3’ single-strands. The Mre11-Rad50-Nbs1 (MRN) complex is a critical component of this pathway as it performs the initial short-range resection events and also facilitates loading and activation of other nucleases that perform long-range processing^4–8^. The MRN complex, together with phosphorylated CtIP protein as an activator, performs this initial processing in the form of endonucleolytic cuts on the 5ʹ strand^6,8^. We and others have previously shown that efficient endonuclease cutting by Mre11 requires the Nbs1 protein, ATP hydrolysis by Rad50, and phosphorylated CtIP^6–8^. In addition, end processing by MRN (as well as the yeast MRX complex) requires a physical block on a DNA end which guides the Mre11 incisions to sites adjacent to the block^8–11^.

During meiosis, it is clear that this critical protein block is the Spo11 protein—a topoisomerase-related enzyme that creates covalent linkages with DNA during meiotic prophase which require MRN(X) for removal and processing for successful strand exchange^12^. Outside of meiosis when Spo11 is not present, however, it is not clear what constitutes the block. In previous work we have shown in vitro with purified recombinant proteins that the DNA-PK complex can play the role of an MRN-activating protein block, stimulating Mre11 endonucleolytic processing on the 5ʹ and 3ʹ strands adjacent to the end^9^. This collaborative function of DNA-PK in DNA end processing is perhaps surprising since the long-held paradigm for DSB repair suggests that NHEJ and replication-dependent pathways act in opposition to each other^13,14^. The widely used term “pathway choice” also has generally been used to describe a competitive relationship between NHEJ and homologous recombination and has implied a competition between DNA-PK and MRN for DNA ends.

Here we investigate the initial processing of DSBs by the MRN complex by analyzing DNA-PK-bound cleaved DNA fragments in human cells genome-wide. We find that the production of these processing intermediates is dependent on Mre11 endonuclease activity and occurs at sites of induced DSBs as well as at spontaneous endogenous sites. The pattern of DNA-PK binding and release at sites adjacent to DSBs depends strongly on the kinase activity of DNA-PKcs and correlates very well with the efficiency of binding of homologous recombination factors. Lastly, we use a light-activated Cas9 system for synchronized DSB induction to show that MRN and DNA-PKcs occupy the same ends prior to generation of DNA-PKcs-bound products. These results provide evidence for a sequential (rather than competitive) model of DSB repair in mammalian cells.

## Results

From previous work we know that MRN/CtIP generates nicks on the 5ʹ strand adjacent to a bound DNA-PK complex, and can also generate double-strand breaks through endonucleolytic cleavage of both strands (summarized in Fig. 1A)^9^. To characterize these products of MRN/CtIP-dependent end processing associated with DNA-PK from human cells we used the inducible DivA system where ER-fused AsiSI generates double-strand breaks throughout the genome within 2 to 4 hours of 4-OHT addition^15^. We established a modified ChIP protocol to purify DNA-PK-bound DNA fragments that are released from crosslinked chromatin by performing a <u>g</u>entle lysis, followed by immunoprecipitation with a phospho-S2056 antibody, <u>s</u>ize <u>s</u>election, and purification (GLASS-ChIP)^16^. In previous work, we confirmed that these DNA-PK-bound DNA fragments are produced by human cells after AsiSI induction with 4-OHT using quantitative PCR^9^.

**Figure 1.**
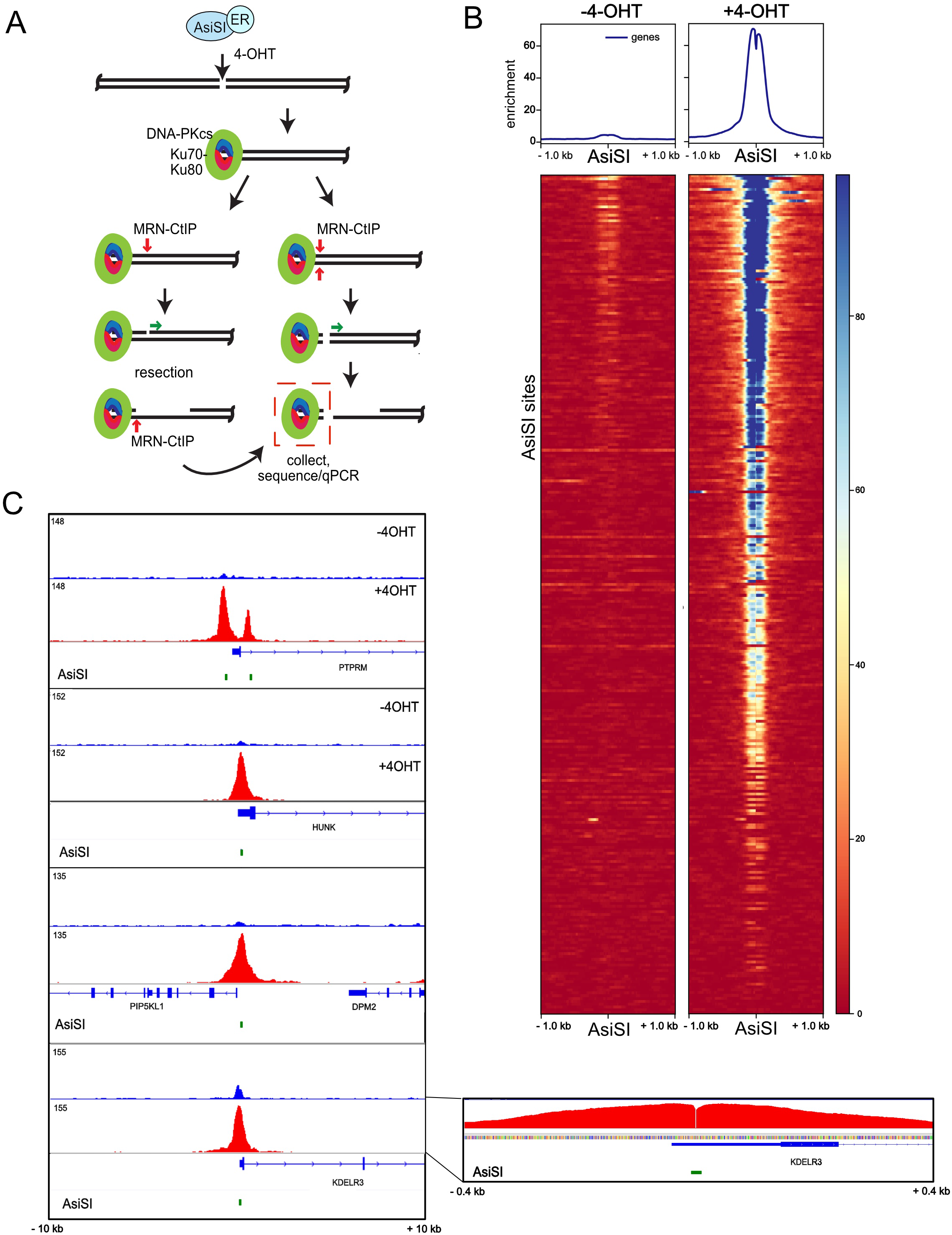
DNA-PKcs-bound fragments are released from DSB sites in human cells. (A) Diagram of DSB induction by ER-fused AsiSI, which translocates into the nucleus with 4-OHT addition. Model for end processing: DNA-PK binds to DSB ends, which are recognized by the MRN complex and CtIP and are processed by endonucleolytic and exonucleolytic nuclease cutting by Mre11. One of the products of this processing is hypothesized to be DNA-PKcs-bound fragments of DNA, released by processing of both strands of DNA simultaneously (red box). These products can be recovered through the GLASS-ChIP procedure and analyzed by sequencing. (B) Recovery of GLASS-ChIP products at the top 300 AsiSI cutting sites is shown, in the absence or presence of 4-OHT as indicated. U2OS cells were induced with 4-OHT for 4 hours in the presence of DNA-PKi (NU7441). (C) Genome browser views of 4 AsiSI sites showing GLASS-ChIP recovery with 4-OHT addition. Inset: high resolution view of one GLASS-ChIP peak showing the absence of signal at the center of the AsiSI cut site.

Here we employed the GLASS-ChIP protocol with human U2OS cells expressing AsiSI, exposed to 4-OHT and DNA-PK inhibitor NU-7441 (DNA-PKi) for 4 hours. We used DNA-PKi because we have found that inhibition of DNA-PKcs increases the yield of MRN-cut DNA fragments by several-fold in reconstitution assays with purified proteins and in human cells^9^. The DNA fragments from cells with or without 4-OHT addition were sequenced and GLASS-ChIP signal from the top 300 most efficiently cut AsiSI sites are shown (Fig. 1B). The result shows essentially no signal in the absence of 4-OHT, while a sharp peak surrounding the AsiSI site is seen with 4-OHT in the presence of DNA-PKi. Examples of genome browser views of several individual AsiSI sites are shown in Fig. 1C. The width of the DNA-PKcs peak in the presence of DNA-PKi is approximately 300 to 400 bp, 150 to 200 bp on each side of the AsiSI site that divides the peak (Fig. 1C, inset). We also observed many genomic sites where DNA-PKcs associates in the absence of DNA damage and found that these sites are often coincident with RNA Polymerase II promoters (examples of 4-OHT-independent sites in Fig. S1).

We have previously shown that nicking at sites of DNA-PK bound to DNA ends is Mre11 nuclease-dependent in vitro^9^. We also found in that work that the removal of DNA-PK from the ends of lambda DNA in single-molecule DNA curtains assays did not occur when an Mre11 nuclease-deficient mutant enzyme was used. In human cells, however, it is challenging to eliminate all functional Mre11 activity since even 5% of the endogenous protein can carry out nuclease activity similar to wild-type cells (unpub. observations) and complete removal of Mre11 is cell-lethal^17^. To reduce Mre11 nuclease activity in cells we utilized the endonuclease and exonuclease inhibitors described previously, PFM01 and PFM39, respectively^18^. Exposure of U2OS cells to these inhibitors and DNA-PKi during 4-OHT induction of AsiSI translocation showed that loss of Mre11 endonuclease activity, but not exonuclease activity, substantially reduces the yield of GLASS-ChIP product at AsiSI sites (Fig. 2A, B). We also observed a similar pattern at sites of DNA-PKcs release from non-AsiSI locations (Fig. 2C).

**Figure 2.**
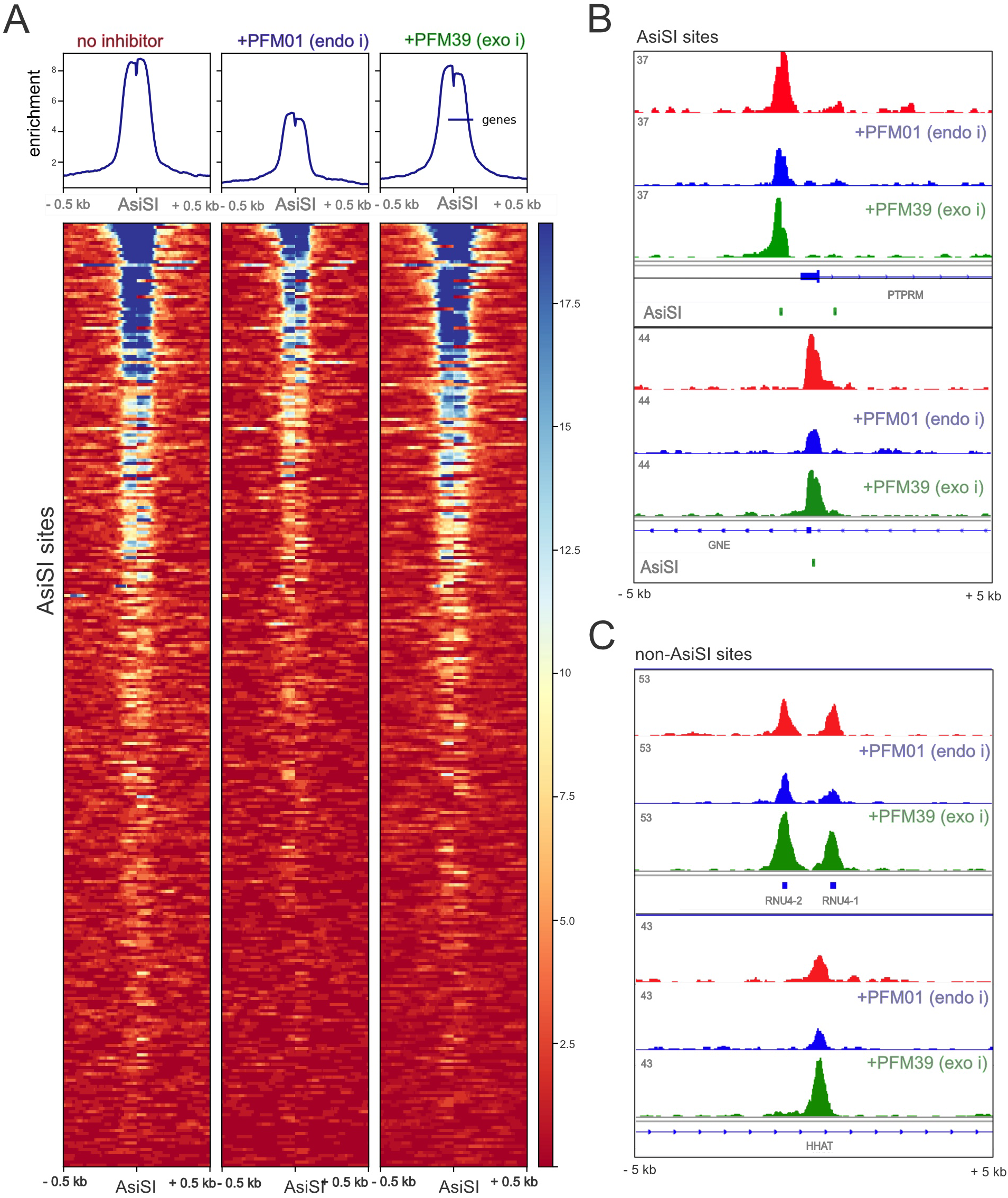
Inhibition of Mre11 nuclease activity blocks release of DNA-PKcs-bound fragments. (A) Recovery of GLASS-ChIP products at the top 300 AsiSI cutting sites is shown in the absence or presence of PFM01 (Mre11 endonuclease inhibitor), or PFM39 (Mre11 exonuclease inhibitor) as indicated. U2OS cells were induced with 4-OHT for 4 hours in the presence of DNA-PKi (NU7441). (B) Genome browser views of 2 AsiSI sites showing GLASS-ChIP recovery with 4-OHT addition and inhibitors as indicated. (C) Genome browser views of non-AsiSI sites showing GLASS-ChIP recovery with 4-OHT addition and inhibitors as indicated.

### DNA-PK-bound chromatin fragments extend far from the break site

In the absence of DNA-PK inhibitor we previously observed 4-OHT-dependent GLASS ChIP recovery by qPCR, although the levels were several-fold lower than what is observed with DNA-PKi present^9^. Here we sequenced GLASS-ChIP libraries made from cells in the absence of inhibitor and found substantial recovery of 4-OHT-induced fragments, although the fold increase over background levels in the absence of 4-OHT varies substantially depending on the site (Fig. 3A). Aggregated data from the top 300 AsiSI sites is shown in Figure 3B, with data from individual genes in Fig. 3C. One obvious difference between inhibitor-free samples and DNA-PKi-treated samples is the fact that DNA-PKcs ChIP signal extends much farther from the AsiSI break site in the absence of DNA-PK inhibition. The recovered fragments cover at least 1 kb from the AsiSI site, in some cases up to several kb, while signal is limited to less than 200 bp in the presence of DNA-PKi.

**Figure 3.**
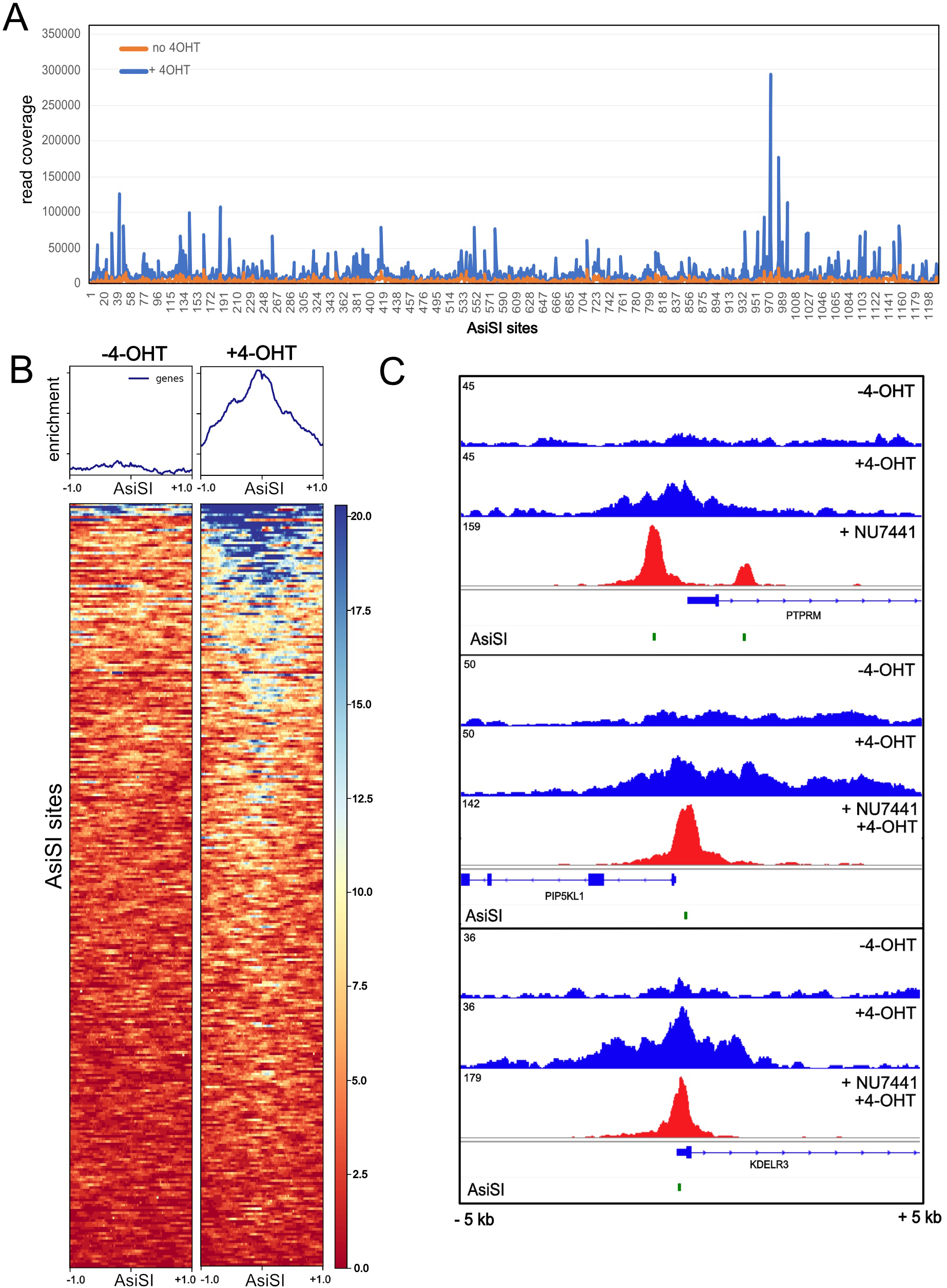
Recovery of DNA-PKcs-bound fragments occurs in the absence of DNA-PKi. (A) Summary of GLASS-ChIP enrichment (total read depth) at all 1211 AsiSI sites (includes 1 kb region upstream and downstream of site), in the absence of DNA-PKi. (B) Recovery of GLASS-ChIP products at the top 300 AsiSI cutting sites is shown, in the absence or presence of 4-OHT (4 hr) as indicated. (C) Genome browser views of 3 AsiSI sites showing GLASS-ChIP recovery with 4-OHT addition and comparison with samples exposed to DNA-PKi.

The vast majority of GLASS-ChIP peaks in the genome are not dependent on 4-OHT exposure (examples shown in Fig. S2). In general, these DNA-PK ChIP signals are significantly higher than AsiSI-associated peaks and are tightly correlated with RNA Polymerase II, both the total polymerase and phosphorylated S2-RNAPII ChIP (RNAP2S2), based on comparisons of previously published ChIP data in U2OS cells^19^. This observation agrees with many reports of DNA-PKcs association with sites of active transcription and even a requirement for DNA-PK in transcriptional regulation in some cases^20–23^. These sites are often not associated with DSBs, as shown by a lack of BLESS signal, and also no accumulation of other DSB markers such as Lig4, 53BP1, and Rad51^24^ (Fig. S2A). In other cases, the DNA-PKcs GLASS-ChIP signal is found at sites that also show several other markers of DNA double-strand breaks, including Lig4, 53BP1, and BLESS signal (examples of both types of sites in Fig. S2B).

To compare the yield of DNA-PKcs-bound fragments released by end processing to the pattern of DNA-PKcs occupancy in the genome, we performed conventional DNA-PKcs ChIP using phospho-DNA-PKcs antibody with extensive sonication of chromatin (DNA-PKcs “Pellet-ChIP”) in the absence of DNA-PKi. Recovery of DNA-PKcs-bound pellet ChIP fragments at the top 300 AsiSI sites appears to be qualitatively very similar to what is recovered from the released GLASS-ChIP fragments (Fig. 4A), and a comparison to the GLASS-ChIP recovery of fragments at these 300 sites shows a ρ correlation coefficient of 0.79 (Fig. 4B). Genome browser views of the released versus chromatin-bound DNA-PKcs show that the patterns are generally very similar, with many sites independent of 4-OHT as well as AsiSI-associated peaks (arrows, + 4-OHT, Fig. 4C). Higher resolution views show that, in some cases, the spreading of DNA-PKcs away from the AsiSI site is more pronounced with the GLASS-ChIP pattern (Fig. 4C panels 2 and 4). Consistent with these observations, the average width of the GLASS-ChIP peak is approximately 850 bp on either side of AsiSI, while the corresponding width in the pellet-ChIP samples is approximately 700 bp.

**Figure 4.**
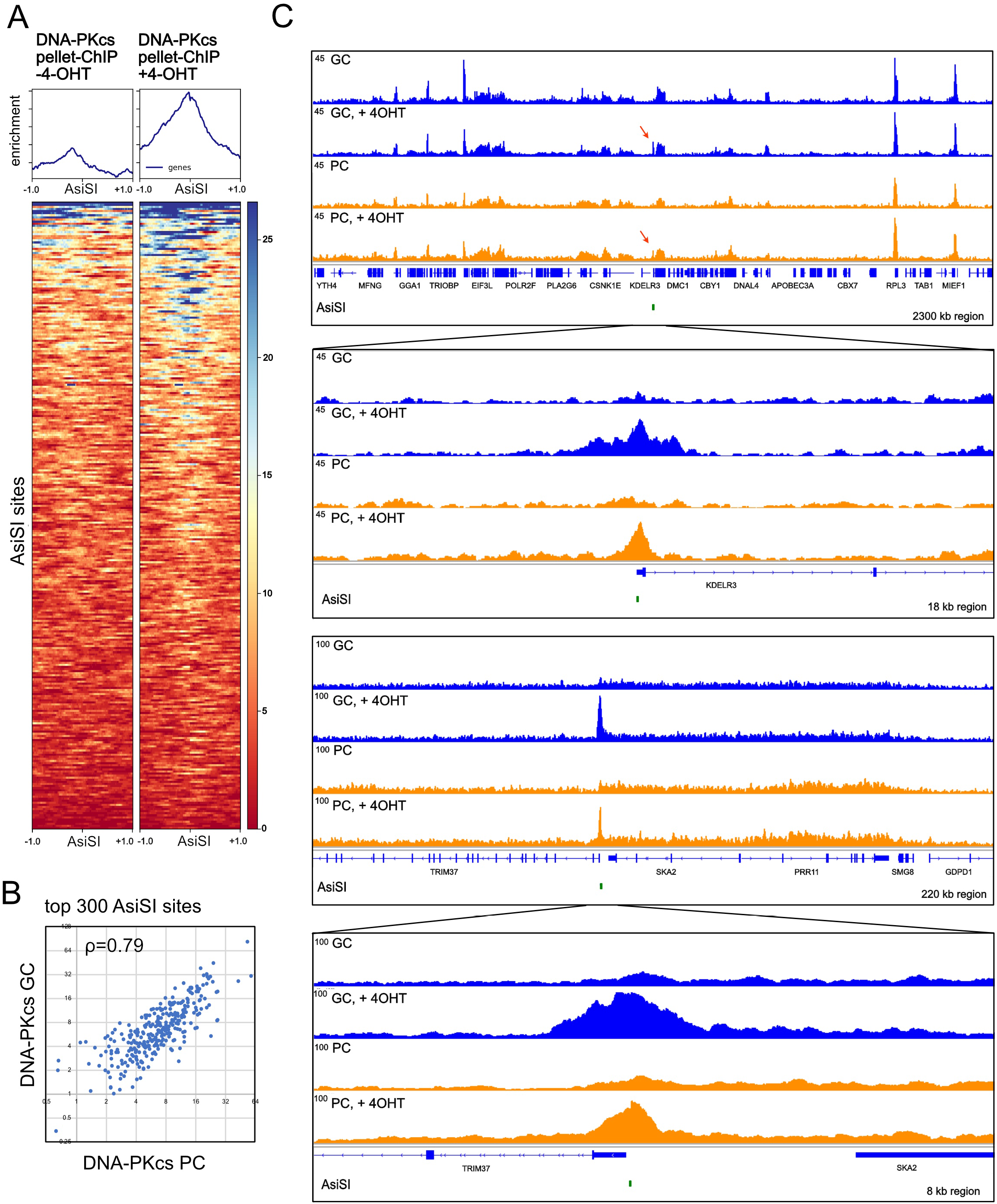
Comparisons between DNA-PKcs released and chromatin-bound fragments in the absence of DNA-PKcs inhibition. (A) Recovery of DNA-PKcs pellet-ChIP products at the top 300 AsiSI cutting sites is shown, in the absence or presence of 4-OHT (4 hrs) as indicated. (B) enrichment for DNA-PKcs GLASS-ChIP versus pellet ChIP is shown for the 300 most efficiently cut AsiSI sites, with Pearson correlation coefficient as indicated. (C) Genome browser views of 2 AsiSI sites showing GLASS-ChIP and DNA-PKcs pellet-ChIP recovery with 4-OHT addition. Insets show higher resolution views.

### DNA-PKcs-bound GLASS-ChIP product efficiency aligns with Rad51 as well as NHEJ factors

To quantitatively assess sites throughout the genome for DNA-PKcs occupancy and other markers, the top 2500 DNA-PKcs-enriched genomic sites from the GLASS-ChIP dataset were analyzed for previously reported chromatin marks as well as for DSB-binding factors^19,20,24^. Binding for each factor or mark was ranked within the 2500 genomic locations and analyzed by unsupervised hierarchical clustering (Fig. S3). This analysis shows that approximately 20% of the highest efficiency DNA-PKcs-bound sites in U2OS cells are also associated with binding of DSB factors Lig4, Xrcc4, 53BP1, and Rad51, likely indicating the presence of naturally-occurring DSBs. Other chromatin marks are also closely associated with a subset of these sites, including γ-H2AX, H3K36me2, and H4S1p. The remaining 70% to 80% of the highest DNA-PKcs binding sites are associated with high levels of RNAPII but not with DSB-associated marks.

Looking more closely at the AsiSI-induced break sites, we analyzed the efficiency of GLASS-ChIP or DNA-PKcs pellet-ChIP at the top 300 restriction enzyme sites and their correlation with DSB-binding factor binding and chromatin marks (Fig. 5). Interestingly, the binding with highest^25^ correlation observed with GLASS-ChIP efficiency is Rad51, followed by the DSB factors Xrcc4, Lig4, and RNAPIIpS2 (Fig. 5A). The pellet-ChIP signal also aligned closely with these DSB-associated factors, although less so compared to the released DNA-PKcs signal, and the DNA-PKcs pellet ChIP showed a much higher correlation with specific marks associated with an open chromatin state and transcription activation (H4K20me1, H2BK120Ub, and H3K36me3)^26,27,27,28^. Consistent with the close association between the efficiency of DNA-PKcs ChIP recovery and DSB-associated factors, we found that hierarchical clustering of the AsiSI genomic locations for all of the reported chromatin marks shows that DNA-PKcs (both GLASS-ChIP and pellet-ChIP) clusters together with Rad51, 53BP1, Lig4, and Xrcc4 (Fig. 5B).

**Figure 5.**
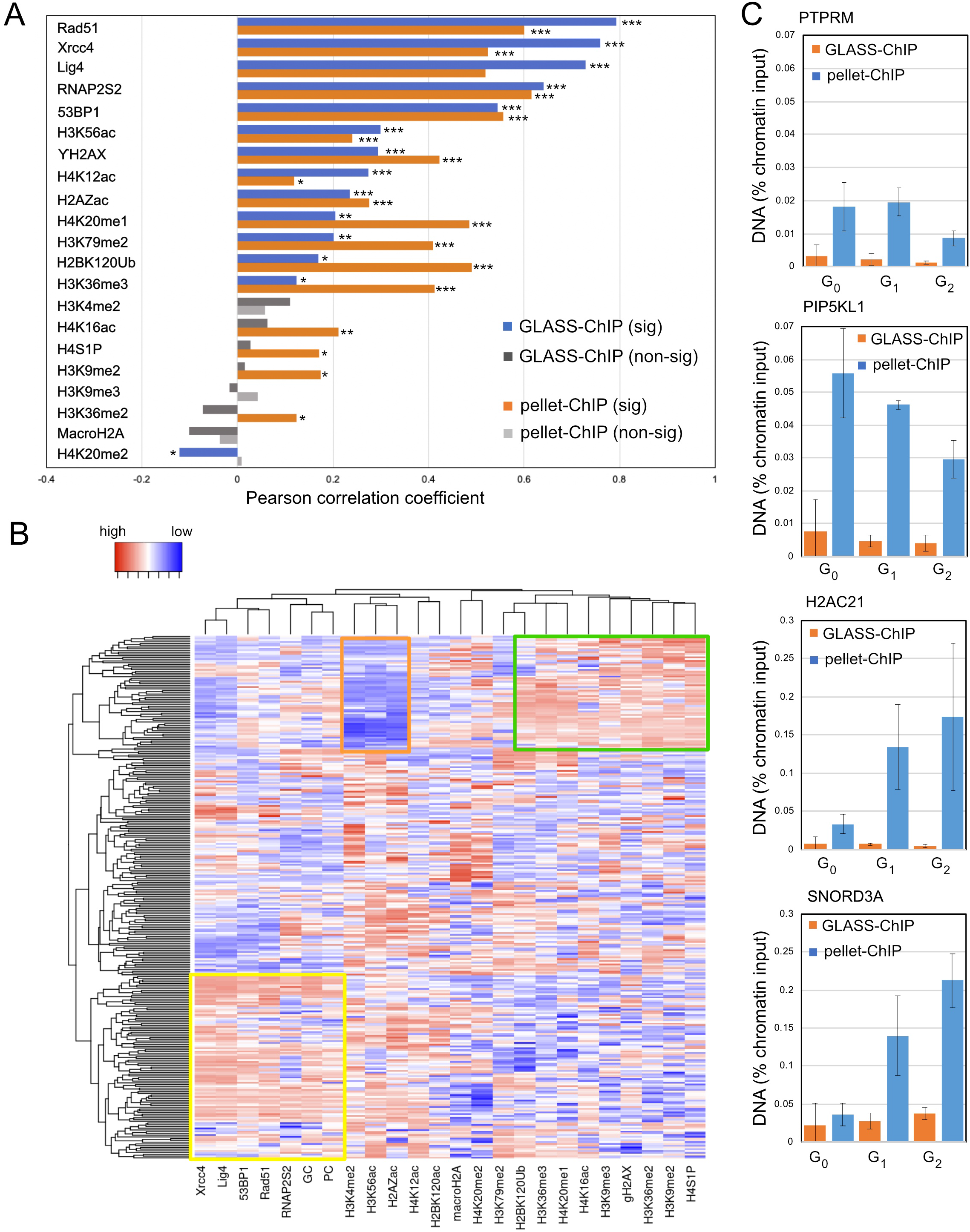
DNA-PKcs released fragments and pellet-ChIP efficiency correlate with DSB markers. (A) Enrichment for DNA-PKcs in GLASS-ChIP and DNA-PKcs pellet-ChIP datasets in the absence of DNA-PKcs inhibition was compared with published data showing enrichment for DSB-associated binding factors and chromatin marks^19,20,24^, generating correlation coefficients as shown. p values calculated using the linear regression t test; *, **, *** indicate p<0.05, 0.0005, and 0.00005, respectively. (B) Heat map showing unsupervised clustering of DNA-PKcs GLASS-ChIP (GC), pellet-ChIP (PC) and enrichment datasets showing DSB-associated binding factors and chromatin marks^19,20,24^ using ranked enrichment data at the top 300 AsiSI sites. Boxes indicate clusters of breaks with DSB-associated marks (yellow), H3 methylation associated marks (green), and low H3K56ac and associated marks (orange). (C) Recovery of DNA-PKcs GLASS-ChIP and pellet-ChIP at two AsiSI sites (PTPRM, PIP5KL1) and two non-AsiSI genomic locations (H2AC21 and SNORD3A) by quantitative PCR. Error bars indicate standard deviation. Results are shown relative to chromatin input.

Some of the chromatin marks showing significant correlations with DNA-PKcs ChIP recovery are associated with S phase or mitosis^29,30^ and thus may be correlated due to cell cycle specificity. To determine if the MRN cutting adjacent to DNA-PKcs is specific to certain cell cycle phases, we tested the efficiency of GLASS-ChIP in G_0_, G_1_, and G_2_ phases by qPCR at two AsiSI sites PTPRM and PIP5KL1 (Fig. 5C). This analysis showed that GLASS-ChIP products are observed in all cell cycle phases at approximately 10% efficiency compared to DNA-PKcs CHIP from the genome (pellet-ChIP). A similar analysis with non-AsiSI DNA-PKcs binding sites H2AC21 and SNORD3A shows varying ratios depending on the gene, while the levels of DNA-PKcs bound at these sites increases substantially in G_1_ and G_2_ phases relative to G_0_. This may be related to increased transcription of these genes since histone biosynthesis is cell cycle-regulated^31^ and small nucleolar RNAs are tied to ribosomal RNA maturation which is also determined by cell cycle phase^32^. To confirm that resection of AsiSI-induced breaks actually occurs in all cell cycle phases, we measured resection using a qPCR method^25^ (Fig. S4). These results confirmed that a low level of resection occurs in G_0_ cells as measured by this assay, but is significantly higher in G_1_ and G_2_ cells. We also confirmed that exposure to DNA-PKi increases resection efficiency as we previously reported^33^.

### DNA-PKcs and Mre11 coincide at DSB sites before MRN cleavage product is formed

Although we have documented the binding and release of DNA-PKcs from chromatin with AsiSI induction, it is challenging to characterize the kinetics of these events with this system since the tamoxifen-induced translocation of estrogen receptor fusions occurs gradually over the course of several hours and the AsiSI enzyme also has the opportunity to cut religated sites multiple times^34^. Thus we turned to the recently developed Cas9-based system where a guide RNA containing caged nucleotides allows for Cas9 recognition of a target site, but does not undergo cleavage of the DNA until cells are exposed to light of an appropriate wavelength to remove the caged adduct (365 nm)^35^. This CRISPR system allows for high temporal resolution of events since Cas9 cleavage of the target site occurs within seconds of light exposure. Subsequent work using this light-activated system also generated several multi-site guides, each of which recognizes a large number of identical targets in the genome^36^. Here we used one of these multi-target guides (AluGG) that recognizes 117 copies of a short interspersed nuclear element.

With the light-activated Cas9 in HEK293T cells, we performed GLASS-ChIP of DNA-PKcs released from chromatin, in the presence of DNA-PKi (Fig. 6A). Analysis of all 117 target sites shows robust product accumulation at 60 min. post light exposure, but no DNA-PKcs fragments at 15 min. (Fig. 6B). In contrast, Mre11 ChIP performed under the same conditions shows accumulation of MRN at the 15 min. time point (Fig. 6C), as reported previously in the absence of DNA-PKi^36^. Thus, the MRN complex is present at the DSB early after break induction but the DNA-PKcs product requires a longer occupancy at the break site.

**Figure 6.**
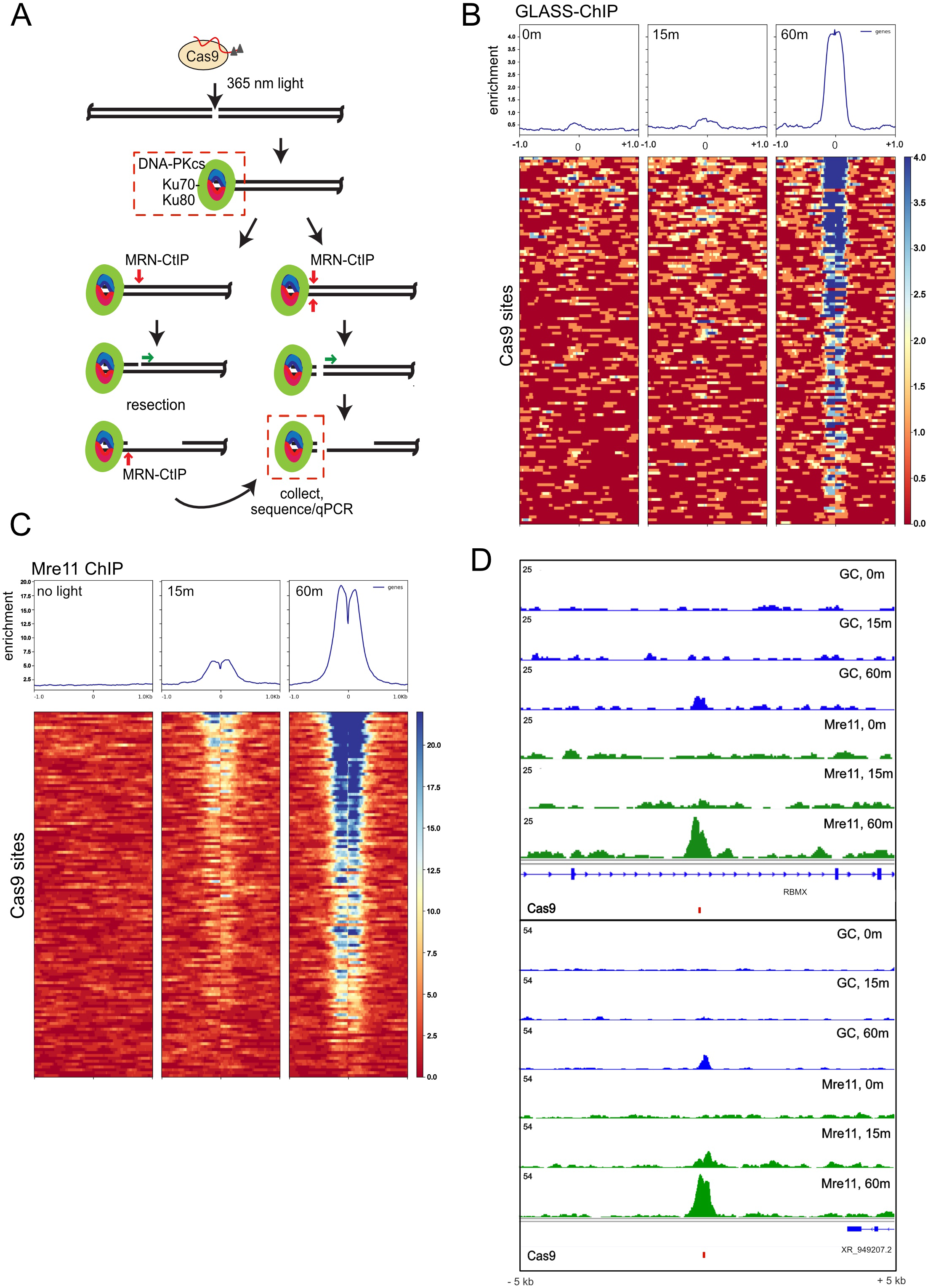
GLASS-ChIP recovery of DNA-PKcs-bound fragments at light-activated Cas9 target sites. (A) Diagram of DSB induction by light-activated Cas9. Model for end processing as in Fig. 1A. (B) Recovery of GLASS-ChIP DNA-PKcs products at the 117 target sites for the AluGG guide RNA are shown, without 365 nm light activation (0 min) or 15 or 60 min. following light exposure as indicated, in the presence of DNA-PKi (NU7441). (C) Recovery of Mre11 ChIP products at the 117 target sites for the AluGG guide RNA are shown, without 365 nm light activation (0 min) or 15 or 60 min. following light exposure as indicated, in the presence of DNA-PKi (NU7441). (D) Genome browser views of 2 Cas9 target sites showing GLASS-ChIP and Mre11 recovery at 0, 15 min., or 60 min. post light exposure.

We also examined the DNA-PKcs present in the chromatin at varying time points after light exposure and found that the protein is present at the target sites at an early time point (15 min., DNA-PKcs pellet-ChIP)(Fig. 7A). Comparison of Mre11 and DNA-PKcs pellet ChIP patterns show a similar accumulation at this time whereas the released GLASS-ChIP product is not present (Fig. 7B), suggesting that the MRN complex and DNA-PKcs occupy the same sites immediately following break induction and that the DNA-PKcs-bound product is released subsequent to this co-localization. Examination of the locations of ChIP products at the 15 min. time point suggests that the chromatin-bound DNA-PKcs (pellet-ChIP, PC) is located at the break site while Mre11 is positioned adjacent to this, away from the break site (Fig. 7C). Genome browser views of two Cas9 target sites show examples of this coincident accumulation of DNA-PKcs and Mre11 at specific cut sites (Fig. 7D).

**Figure 7.**
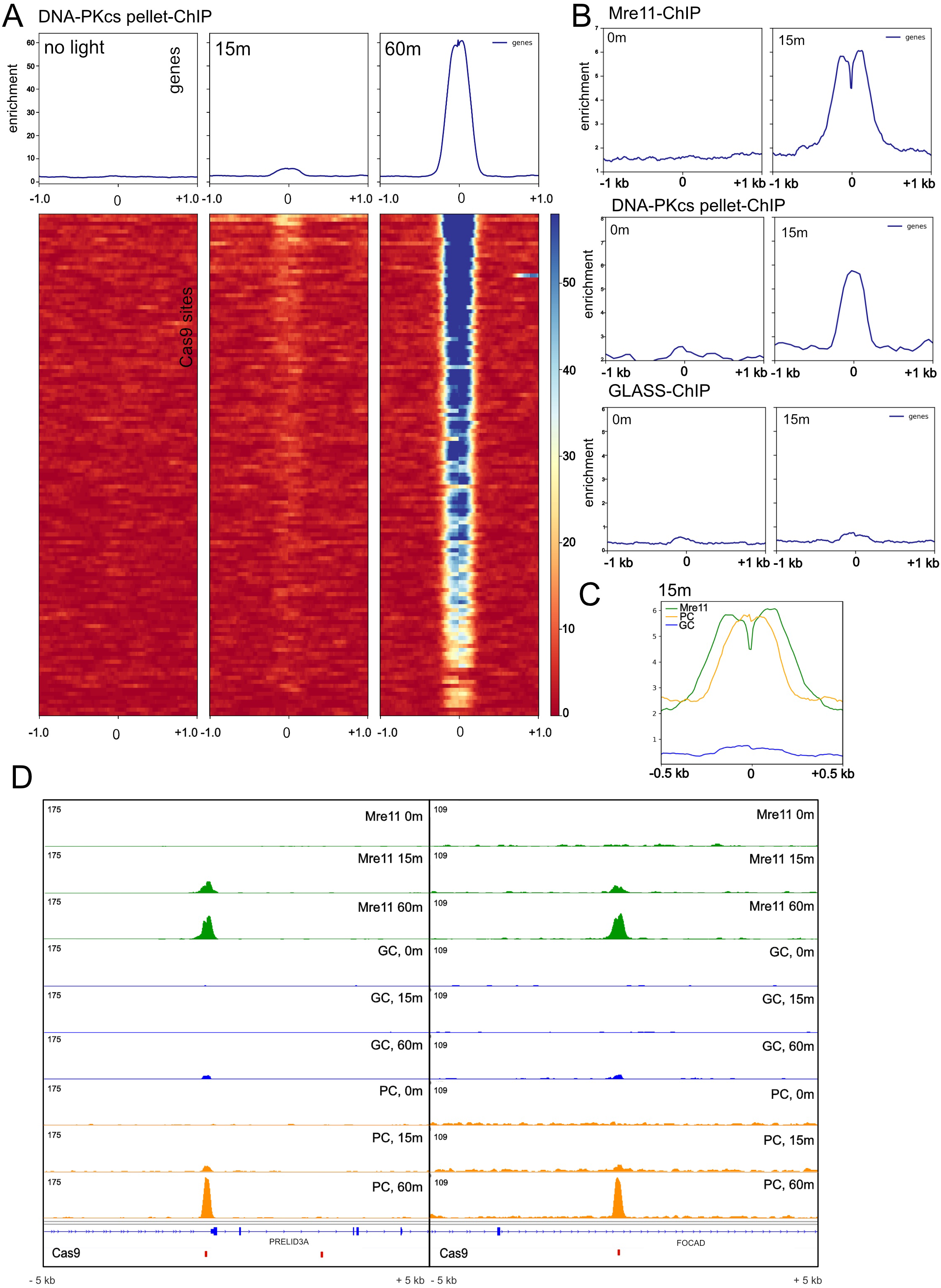
DNA-PKcs and Mre11 bind to DSB site before DNA-PKcs is released. (A) Recovery of DNA-PKcs ChIP products (pellet ChIP) at the 117 target sites for the AluGG guide RNA are shown, without 365 nm light activation (0 min) or 15 or 60 min. following light exposure as indicated, in the presence of DNA-PKi (NU7441). (B) Comparison of Mre11 ChIP, DNA-PKcs pellet ChIP, and GLASS-ChIP products at the 0 min. and 15 min. time points, using equivalent enrichment scales. (C) View of Mre11 ChIP, DNA-PKcs pellet ChIP (PC), and DNA-PKcs GLASS-ChIP (GC) recovery at the 117 target sites at 15 min. (D) Genome browser views of 2 Cas9 target sites showing GLASS-ChIP, DNA-PKcs pellet-ChIP, and Mre11 ChIP at 0, 15 min., or 60 min. post light exposure.

## Discussion

In this study we document the production of MRN-dependent fragments of DNA, bound by DNA-PK, and released from the genome at sites of DNA DSBs. Production of these fragments is sensitive to the endonuclease-specific Mre11 inhibitor PFM01. This result, combined with data from purified protein reconstitution experiments and single-molecule assays^9^, suggests that MRN endonuclease activity processes DNA-PK-bound ends in genomic DNA to release short, double-stranded products.

From the in vitro assays we and others have performed^6,8–11,37^, we know that a protein block is essential for MRN(X) endonucleolytic cutting at DSBs. DNA-PK is an excellent candidate for such a block because it binds tightly to ends through the Ku heterodimer. We also know from laser irradiation studies that it is one of the first complexes to arrive at a DSB, generally faster than Mre11 kinetics^38–40^. In vitro we observed both 5ʹ nicks as well as concurrent 5ʹ and 3ʹ cutting by MRN, which generates a new DSB^8,9^, with the nicking activity about 7 to 8-fold more efficient than double-strand cuts. The position of the endonuclease cleavage in vitro with DNA-PK, MRN, and CtIP present is approximately 45 nt from the end of the 5ʹ strand. In cells, however, with DNA-PKi present, we find that the fragments are significantly larger—on average 150-180 nt from the AsiSI cut site. We do not know the reasons for this difference in cut location but speculate that other end-binding factors present in cells that associate with DNA-PK likely contribute to the size of the end-bound complex. Mass spectrometry characterization of proteins bound to the DNA-PK-associated fragments show the MRN complex and several other DNA repair factors including PARP, Xrcc1, FEN1, and Mdc1. In addition, a large number of RNA splicing and transcription-associated proteins are enriched on these ends (Fig. S5).

When DNA-PK is inhibited in vitro, the complex remains bound very stably to DNA ends^41^. This nearly irreversible binding is due to the fact that DNA-PKcs autophosphorylation is required to promote release from DNA^42–44^ and that the phosphorylation of Ku70 by DNA-PKcs also promotes release of Ku^45^, so when these events are blocked, DNA-PK is immobilized. In vitro, DNA-PKi strongly promotes MRN nuclease activity^9^, similar to the robust release of DNA-PK-bound fragments in the presence of DNA-PKi observed here in cells.

In the absence of DNA-PKi, induction of DSBs by AsiSI produces DNA-PK-bound fragments that map to sites directly adjacent to the DSB location but also appear farther away— up to 1 kb or more depending on the site. It is not clear what generates this distribution. One possibility is that multiple binding and cutting events occur over the time period of AsiSI translocation into the nucleus. Alternatively, there could be movement of Ku on DNA such that sliding and stacking of DNA-PK occurs, ultimately generating released products that extend far from the original site of the DSB. We favor the latter model, based on early experiments with DNA-PK in human cell nuclear extracts that suggested movement of the complex inward from DNA ends in a manner dependent on DNA-PK catalytic activity^46^. This contrasts with the ATP-independent sliding of Ku alone^47,48^.

Interestingly, the extent of DNA-PKcs spreading observed using GLASS-ChIPseq in the absence of DNA-PKi is more extensive than observed in standard pellet-ChIP using the same conditions. Interpreted in light of the sliding model, this result would suggest that MRN processing of DNA-PK-bound fragments occurs more efficiently at sites of internalized DNA-PK compared to all DNA-PK-bound sites.

With the AsiSI-ER DivA system, characterization of the kinetics of DNA-PK removal is challenging due to the extended period of AsiSI translocation into the nucleus. The light-activated Cas9 system^35,36^, however, allows for precise control of the cleavage step and therefore gives us the opportunity to look at early time points following break induction. Here we found that Mre11 and DNA-PK are both found on chromatin adjacent to the Cas9 cut site at an early time following light exposure (15 min.). At this time, released DNA-PK-bound fragments (GLASS-ChIP) are not observed, but production of these fragments is clearly seen at a later time point (60 min). Thus, we infer that MRN and DNA-PK likely occupy the same ends for some time after break induction (overlapping with the 15 min. time point) and that MRN processing of these ends occurs after this simultaneous binding period. From the pattern of ChIP occupancy we observed, DNA-PK is bound at the break site while MRN is located adjacent to this, away from the end. This interpretation is also supported by our previous single-molecule observations where MRN colocalization with DNA-PK on DNA ends was frequently observed and preceded Mre11 nuclease-dependent loss of both DNA-PK and MRN from the ends^9^. In these experiments we also observed MRN complexes internal to DNA-PK-bound ends before release. We do not know what the trigger is for the processing event, but it is likely a conformational change inherent to the MRN, CtIP, and DNA-PK complexes considering that the in vitro experiments utilized purified recombinant proteins.

The recovery of DNA-PKcs-bound DNA fragments genome-wide provides a global view of binding preferences. Correlation analysis of DNA-PKcs binding efficiency with other DNA repair proteins, chromatin marks, and transcription-related factors shows that the binding is most tightly correlated with RNAPII. DNA-PKcs has been observed to associate physically and functionally with RNAPII previously and in some cases was observed to be essential for transcription activation^22,23,49,50^. The exact role of DNA-PK in this context, however, has not been defined. Here we see that approximately 20% of the most efficiently bound DNA-PK sites in the genome that are coincident with RNAPII appear to be DSB sites, on the basis of coincident Xrcc4, Lig4, and 53BP1 signals. Considering that DSB formation has been shown to be essential for transcription in some contexts, it is possible that the DSB machinery is regulating transcription at these genomic locations. The remainder of the DNA-PKcs genomic binding sites show RNAPII but are devoid of other DSB factors. Recent work also shows that MRN occupancy in the genome is strongly correlated with RNAPII abundance^51^.

Correlation analysis of DNA-PKcs binding in comparison with previously published U2OS DivA datasets from the Legube laboratory also shows that GLASS-ChIP recovery within the most efficiently cut AsiSI sites shows the highest correlation with Rad51 and other DSB markers, including Xrcc4, Lig4, and 53BP1. This is consistent with expectations, considering that MRN processing is required for subsequent long-range resection and Rad51 loading. Hierarchical clustering of all of the AsiSI sites shows these preferences clearly, with the most efficiently cut sites clustering together and binding all of the DSB-specific factors. From this analysis there do not appear to be separate groups of DSBs that bind to Rad51 or the NHEJ-related factors exclusively but all of these factors associate with the sites that are the most efficiently cut.

From our results in this work as well as our previous in vitro studies, we propose that DNA-PK plays an essential role in DSB processing by acting as a protein block to stimulate MRN end processing (Fig. 8). Most of this processing is likely in the form of 5ʹ strand nicks, based on ensemble reconstitution assays with purified proteins, while approximately 10 to 15% of the events lead to double-strand processing, releasing a DNA-PK-bound fragment. Overall this presents a very different view of “pathway choice” compared to the canonical model by positing that this must be a sequential series of events, initiated by DNA-PK first binding to DNA ends. Homologous recombination factors such as MRN are thus dependent on and collaborating with NHEJ factors, an “entwined” rather than competitive scenario also predicted by mechanistic modeling of previously published data^52^. In future work it will be important to define the rate-limiting step(s) for Mre11 cleavage and to better understand the role of DNA-PK internalization in promotion of MRN activity to elucidate the details of this collaboration with greater resolution.

**Figure 8.**
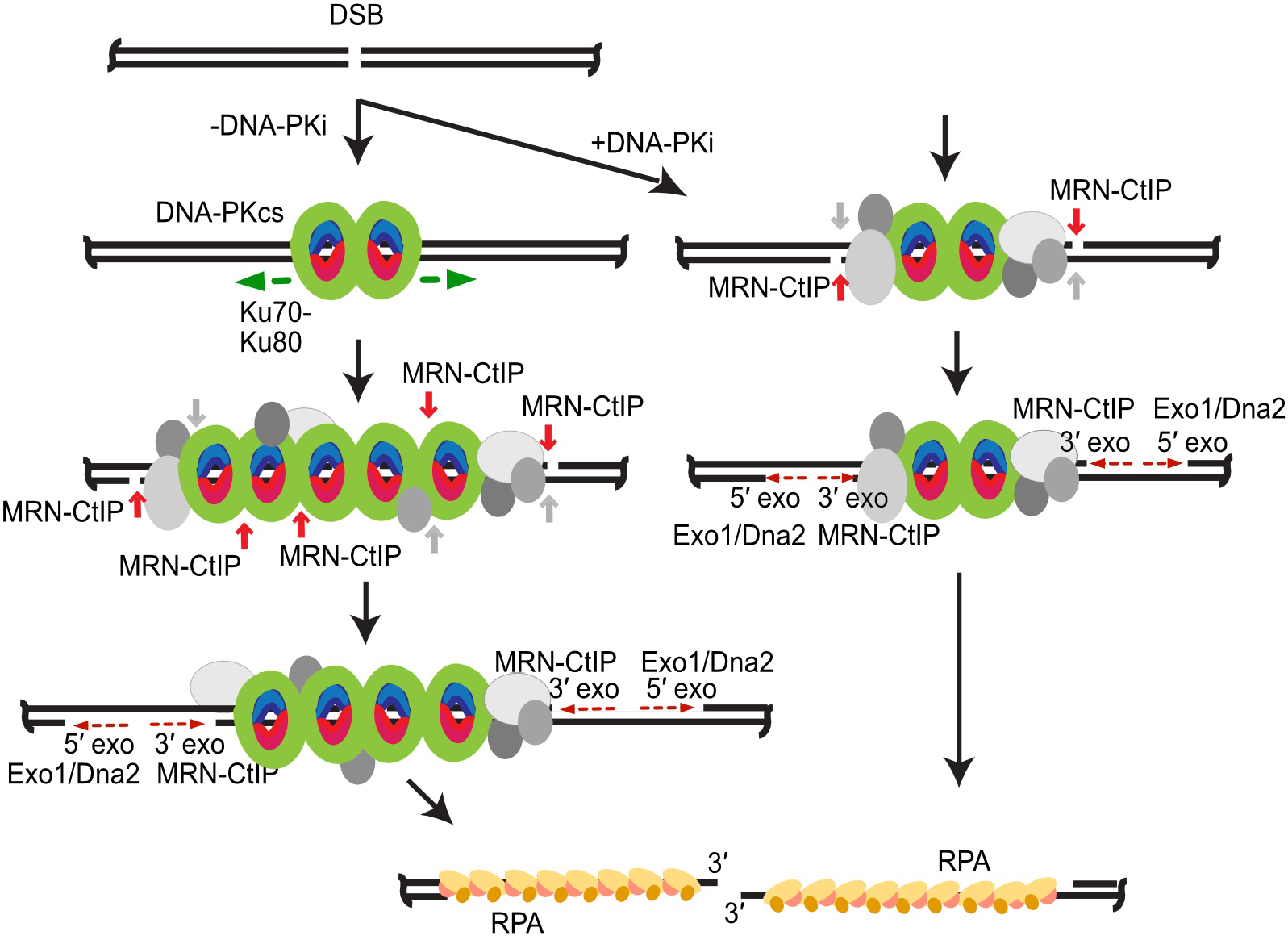
Working model for DNA-PK and MRN dynamics at DSB sites. After DSB formation, DNA-PK (DNA-PKcs green, Ku red/blue) binds to DSBs. In the presence of DNA-PKi (right), DNA-PK remains tightly bound at the DSB site, together with accessory proteins (grey). MRN nicks on the 5′ strands occur (red) and, less frequently, simultaneous nicks on the 3′ strands occur (grey). Simultaneous nicks on both strands give rise to GLASS-ChIP products. Further processing on the 5′ strands by MRN (3′ to 5′ exonuclease, red dashed arrow towards DSB) and by Exo1 or Dna2 (5′ to 3′ exonuclease, red dashed arrow away from DSB) extend nicks into gaps of varying length. Ultimately, gapped intermediates are bound by RPA (yellow/orange) and DNA-PK is removed. In the absence of DNA-PKi (left), DNA-PK slides inward (green arrows), allowing more DNA-PK molecules as well as accessory factors to bind. Alternative model: multiple rounds of cutting and re-binding allow for DNA-PK movement inward (not shown). MRN cutting on 5′ strands and less efficient cutting on 3′ strands occurs as described above, followed by exonucleolytic expansion of the gap.

## Acknowledgments

We thank members of the Paull laboratory for helpful discussion and acknowledge NIH grant R01GM138548 for support. DNA sequencing of DNA-PKcs GLASS-ChIP and pellet-ChIP products was performed at the Genomic Sequencing and Analysis Facility at UT Austin.

## Author contributions

R.D. contributed to conceptualization, methodology, validation, formal analysis, investigation, visualization, and review and editing of the manuscript. A.M-G. contributed to methodology, vlidation, formal analysis, investigation, and review and editing of the manuscript. T.H. contributed to methodology, resources, supervision, and review and editing of the manuscript. T.P. contributed to conceptualization, methodology, validation, formal analysis, investigation, resources, data curation, writing and editing of the manuscript, visualization, supervision, project administration, and funding acquisition.

## Materials and Methods

### Cell Culture

Human U2OS cells were grown in DMEM with high glucose, L-glutamine, and sodium pyruvate and 10% fetal bovine serum, with 1X PEN/STREP. Recombinant AsiSI was expressed via the DivA system as previously described^15^.

### GLASS-ChIP assay

Human U2OS cells with inducible AsiSI^15^ were grown to 50-60% confluency in 150 mm dishes and treated with 600 nM 4-OHT for 4 h at 37°C. When used, 10 *µ*M NU7441 was added 30 min. prior to 4-OHT addition. After 4-OHT treatment, cells were fixed with 1% formaldehyde for 7 min at RT with gentle rotation. Crosslinking was stopped by addition of 125 mM glycine for 5 min, washed twice with cold PBS, harvested, flash frozen in liquid nitrogen and stored at -80°C. For the GLASS-ChIP assay, we modified a standard protocol (Abcam) with a gentle lysis procedure and minimal, low-level sonication to rupture cells without extensive DNA damage as previously described^16^. The formaldehyde fixed cells were thawed at RT for 5 min, resuspended in RIPA buffer (50 mM Tris-HCl pH8.0, 150 mM NaCl, 2 mM EDTA pH8.0, 1% NP-40, 0.5% Sodium Deoxycholate, 0.1% SDS) with 1x protease inhibitors (Pierce #A32955) and sonicated using a Cell Ruptor at low power, for 10 sec followed by 10 pulses after a 20 sec interval. Cell lysates were then centrifuged at 3000 rpm for 3 min at RT to remove the bulk of chromatin. The supernatant was then incubated with 1.6 *µ*g of anti-DNA-PKcs pS2056 antibodies (abcam 124918) overnight at 4°C, followed by incubation with 25 *µ*l Protein A/G magnetic beads (Pierce) at RT for 2 h. Beads were then washed sequentially once in low salt wash buffer (0.1% SDS, 1% Triton X-100, 2 mM EDTA, 20 mM Tris-HCl pH 8.0, 150 mM NaCl), once in high salt wash buffer (0.1% SDS, 1% Triton X-100, 2 mM EDTA, 20 mM Tris-HCl pH 8.0, 500 mM NaCl), once in LiCl wash buffer (0.25 M LiCl, 1% NP-40, 1% Sodium Deoxycholate, 1 mM EDTA, 10 mM Tris-HCl pH 8.0). Beads were then resuspended in TE buffer (10 mM Tris pH 8.0, 0.1 mM EDTA) and transferred to a fresh tube and finally eluted with 100 *µ*l elution buffer (1% SDS, 100mM NaHCO_3_). Crosslinks were reverted for the elutions (65°C for 24 h) and DNA was purified with a Qiagen Nucleotide Clean up kit. DNA fragments <300 bp from ChIP elutions were separated by pulling down larger fragments using paramagnetic Ampure XP beads: 65 *µ*l of Ampure XP beads were added to the uncrosslinked ChIP elution and mixed thoroughly. After 10 min incubation at RT, beads were isolated using a magnet. The supernatant was collected and 25 *µ*l of fresh Ampure XP beads were added and mixed thoroughly. After 10 min at RT, beads were separated and the supernatant was purified using Qiagen Nucleotide Clean up kit. At the final step DNA was eluted in 60 *µ*l of elution buffer (TE). To monitor the Mre11 nuclease dependence, we treated the human U2OS cells expressing inducible AsiSI with 10 *µ*M NU7441, 100 *µ*M PFM01 or PFM39 and 600 nM 4-OHT simultaneously for 1 hour at 37°C before harvesting cells.

### Pellet ChIP

For DNA-PKcs pellet ChIP, the protocol for GLASS-ChIP was followed through the step where chromatin is isolated. The chromatin fraction was then resuspended in 2.2 ml RIPA buffer and was fragmented using a Diagenode Bioruptor on high setting for 30 min. with 10 sec on, 10 sec off. After removal of debris by centrifuging at 800 g for 3 min., the supernatant was incubated with 1.6 *µ*g of anti-DNA-PKcs pS2056 antibodies (abcam 124918) overnight at 4°C, followed by incubation with 25 *µ*l Protein A/G magnetic beads (Pierce) at RT for 2 h. The rest of the procedure was identical to GLASS-ChIP above except that the AMPure bead size selection was not performed.

### Sequencing library preparation

For both GLASS-ChIP and pellet-ChIP sequencing libraries, the eluted DNA was used to make sequencing libraries using the NEBNext Ultra or Ultra II DNA Library Prep Kit for Illumina (NEB) with NEBNext Multiplex dual index primers with 12 amplification cycles and 2 additional AMPure XP clean-up steps at 0.8X. Libraries were sequenced by the UT Genomic Sequencing and Analysis Facility using a NovaSeq SP platform with PE150 runs.

### Data analysis

Raw data was pre-processed with BedTools fastp^53^, mapped with BWA-MEM^54^, duplicates were removed with SAMTools RmDup^55^, and peaks called with MACS2^56^. For Glass-ChIP, no-antibody sample libraries were used as controls whereas for pellet-ChIP, input DNA libraries were used as controls.

Correlation analysis of GLASS-ChIP with previously published datasets^19,20,24^ was performed using deepTools MultiBigWigSummary on MACS2 treatment output files with either a bed file of the top 300 AsiSI sites or with the top 2500 sites identified by MACS2 narrowpeak output, followed by conversion to rankings and hierarchical clustering of rankings for each factor using the Euclidean method with average linkage and visualization with Heatmapper^57^.

### Electroporation and light activation of Cas9 RNP

In order to assemble Cas9-gRNA RNP complex, 2 *µ*L of 100 *µ*M caged AluGG crRNA (Bio-Synthesis)^36^ was mixed with 2 *µ*L of 100 *µ*M tracrRNA (Integrated DNA Technologies) and heated to 95 °C for 3 min in a thermocycler. The cr:tracrRNA was then allowed to cool on benchtop for 5 min. Afterwards, 3 *µ*L of 10 *µ*g/*µ*L (∼66 *µ*M) of purified Cas9 and 8 *µ*L of dialysis buffer (20 mM HEPES pH 7.5, and 500 mM KCl, 20% glycerol) was added to the annealed 4 *µ*L 50 *µ*M cr:tracrRNA for a total of 15 *µ*L, and was thoroughly mixed by pipetting. This solution was incubated for 20 min at room temperature to allow for RNP formation. HEK293T cells were maintained to a confluency of ∼90% prior to electroporation. 12 million cells were trypsinized with 5 min incubation in the incubator. Trypsin was quenched using 1:1 of complete DMEM and cells were then harvested and centrifuged (3 min, 200 *× g*). The supernatant was removed, and cells were washed with 1 mL PBS (resuspend pellet, centrifuge with same settings as above). After the PBS wash, cells were resuspended in 90 *µ*L of nucleofection solution (16.2 *µ*L of Supplement solution mixed with 73.8 *µ*L of SF solution from SF Cell Line 4D-Nucleofector™ X Kit L) (Lonza), transferred to the 15 *µ*L RNP solution; and 2 *µ*L of Cas9 Electroporation Enhancer (Integrated DNA Technologies) was added. The final solution (approximately 125 *µ*L) was gently mixed and transferred to a 100 *µ*L cuvette (Lonza). Electroporation was then performed according to the manufacturer’s instructions on the 4D-Nucleofector™ Core Unit (Lonza) using code CA-189. A total of 400 *µ*L of DMEM complete was used to completely transfer the cells out of the cuvette, before plating to culture wells pre-coated with 1:100 collagen. Cells were incubated with caged Cas9 RNP for 12 h before light activation. NU-7441 DNA-PK inhibitor was added at 10 *µ*M final concentration 1 hr before light exposure. For Cas9 photo-activation, cells were exposed to 1 min of 365 nm light exposure from a handheld blacklight (Amazon https://www.amazon.com/JAXMAN-Ultraviolet-365nm-Detector-Flashlight/dp/B06XW7S1CS/). Typically, 6 flashlights were used at once. Samples were harvested without light exposure (0 min), or 15m and 1 h after light exposure.

### Mre11 ChIP

A minimum of 4 million Tcells were used for each ChIP reaction. For Mre11 ChIP measurements cells were washed once with room temperature PBS, then scrapped off the plate with 10 mL DMEM and transferred to 15 mL falcon tubes. 721 *µ*L of 16% formaldehyde (methanol-free) was added, and the fixation reaction was incubated for 10 min at room temperature with rotation. 750 *µ*L of 2 M glycine was added to quench the formaldehyde, followed by a 3 min incubation with rotation. Cells were spun down at 1,200 *× g* at 4 °C for 3 min, then washed twice with ice-cold PBS. The supernatant was removed, and the crosslinked cell pellet was resuspended in 4 mL lysis buffer LB1 (50 mM HEPES, 140 mM NaCl, 1 mM EDTA, 10% glycerol, 0.5% Igepal CA-630, 0.25% Triton X-100, pH to 7.5 using KOH, add 1x protease inhibitor right before use) for 10 min at 4 °C, then spun down 2,000 *× g* at 4 °C for 3 min. The supernatant was removed and cells were resuspended in 4 mL LB2 (10 mM Tris-HCl pH 8, 200 mM NaCl, 1 mM EDTA, 0.5 mM EGTA, pH to 8.0 using HCl, add 1x protease inhibitor right before use) for 5 min at 4 °C, spun down with the same protocol. The supernatant was removed, and cells were then resuspended in 1.5 mL LB3 (10 mM Tris-HCl pH 8, 100 mM NaCl, 1 mM EDTA, 0.5 mM EGTA, 0.1% Na-Deoxycholate, 0.5% N-lauroylsarcosine, pH to 8.0 using HCl, add 1x protease inhibitor right before use) and transferred to 2 mL tubes. Sonication was performed using a Fisher 150E Sonic Dismembrator with settings: 50% amplitude, 30 s ON, 30 s OFF for 12 min total time. The sonicated sample was spun down (20,000 *× g* at 4 °C for 10 min), and the supernatant was transferred to a 5 mL tube. 1.5 mL of LB3 (with no protease inhibitor) and 300 *µ*L of 10% Triton X-100 were added, and the entire solution was well mixed by gentle inversion. Beads pre-loaded with antibodies were prepared before cell harvesting. 50 *µ*L Protein A beads (Thermo Fisher) were used per IP and transferred to a 2 mL tube on a magnetic stand. Beads were washed twice with blocking buffer BB (0.5% BSA in PBS), then resuspended in 100 *µ*L BB per IP. 3 *µ*L of Mre11 antibody per IP (MRE11 – Novus NB100-142) was added and the mixture placed on rotator for 1-2 h. Right before IP, the 2 mL tube was placed on a magnetic rack and washed 3x with BB, before resuspending in 50 *µ*L BB per IP. 50 *µ*L of beads in BB were transferred to each IP and placed in 4 °C rotator for 6+ hours.

ChIP samples were transferred to a 1.5 mL LoBind tube on a magnetic stand, washed 6x with 1 mL RIPA buffer (50 mM HEPES, 500 mM LiCl, 1 mM EDTA, 1% Igepal CA-630, 0.7% Na-Deoxycholate, pH to 7.5 using KOH), then washed 1x with 1 mL TBE buffer (20 mM Tris-HCl pH 7.5, 150 mM NaCl). The liquid was removed, and the beads containing ChIP’d DNA were eluted in 50 *µ*L elution buffer EB (50 mM Tris-HCl pH 8.0, 10 mM EDTA, 1% SDS) and incubated 65 °C for 6+ hours to remove crosslinks. 40 *µ*L of TE buffer was mixed to dilute the SDS, followed by 2 *µ*L of 20 mg/mL RNaseA (New England BioLabs) for 30 min at 37 °C. 4 *µ*L of 20 mg/mL Proteinase K (New England BioLabs) was added and incubated for 1 h at 55 °C. The genomic DNA was column purified (Qiagen) and eluted in 41 *µ*L nuclease free water.

For preparation of sequencing libraries, end-repair/A-tailing was performed on 17 *µ*L of ChIPed DNA using NEBNext® Ultra™ II End Repair/dA-Tailing Module (New England BioLabs), followed by adapter ligation using T4 DNA Ligase (New England BioLabs). Libraries were amplified with 13 cycles of PCR using single indexed primers.

For MRE11 ChIP-Seq after Cas9 light activation, ChIP’ed DNA samples were pooled, quantified with QuBit (Thermo), Bioanalyzer (Agilent) and qPCR (BioRad), then sequenced on a NextSeq 500 (Illumina) using high-output paired 2×50bp reads. Reads were demultiplexed after sequencing using bcl2fastq. Paired-end reads were aligned to hg38 using bowtie2. Samtools^55^ was used to filter for mapping quality >= 25, remove singleton reads, convert to BAM format, remove potential PCR duplicates, and index reads.

All raw sequencing files as well as processed data (bigwig files) have been deposited to GEO under record GSE218590. AsiSI files are processed with Hg19 genome build; Cas9 light activated data processed with Hg38.

### Resection

Resection of 5ʹ strands at AsiSI breaks was performed using a qPCR-based method as previously described^25^.

### Cell cycle synchronization

Cells were synchronized in G_0_ by serum starvation for 3 days. G_1_ cells were obtained by addition of serum to G_0_ cells for 8 hours followed by addition of 4-OHT for 4 hrs before harvesting for ChIP or resection assays. G_2_ cells were obtained by growing cells in low-dose (2 μg/ml) aphidicolin overnight to synchronize cells at the G_1/_S phase boundary. The aphidicolin was removed, cells were grown for 6 hours followed by addition of 4-OHT for 4 hrs before harvesting for ChIP or resection assays.

### Quantitative PCR

qPCR monitoring of GLASS-ChIP and pellet-ChIP libraries was performed using primers for 4 different AsiSI sites (Table S1). %DNA was calculated using equation 2^(Ct(input) – Ct(test))^ x 100%. DNA values obtained for IP in absence of antibodies were subtracted from the values obtained in presence of antibody and used for plotting the graphs in Fig. 5.

### Mass Spectrometry

GLASS-ChIP was performed as described above, using no antibody samples as controls, in two biological replicates, except that the cells were crosslinked with formaldehyde for only 1 min. Samples eluted from the ProteinA/G beads were processed through a modified Filter-Assisted Sample Preparation protocol as previously described^58^. Protein identification by LC-MS/MS was provided by the University of Texas at Austin Proteomics Facility on an Orbitrap Fusion following previously published procedures^59^. Raw files were analyzed using label-free quantification with Proteome Discoverer 2.2. Results were further refined by two additional methods; first, all proteins were cross-referenced for common contaminants, in which case they were removed from final analysis, and any polypeptides with less than two unique peptides identified were removed from final analysis. The details of workflow for PD 2.2 are available upon request. Proteins identified in both biological replicates with ratio of recovery in +Ab greater than -Ab samples by a value of 5 or higher are reported in Table S2.

## Supplemental Figures

**Figure S1.**
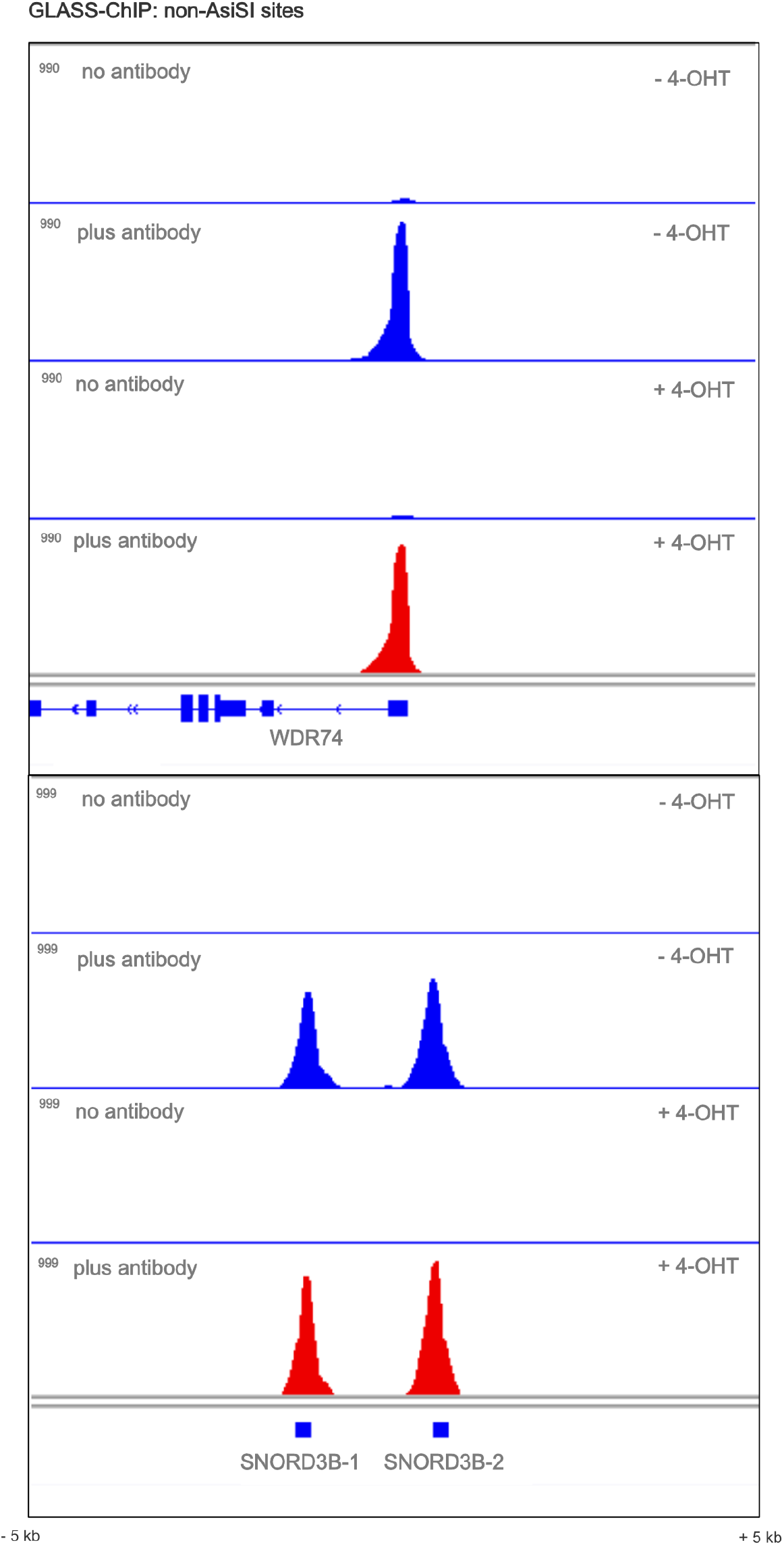
Examples of genome browser views of non-AsiSI sites where phospho-DNA-PKcs is bound in the human genome. No-antibody or Plus-antibody GLASS-ChIP library signals in the absence or presence of 4-OHT are shown as indicated.

**Figure S2.**
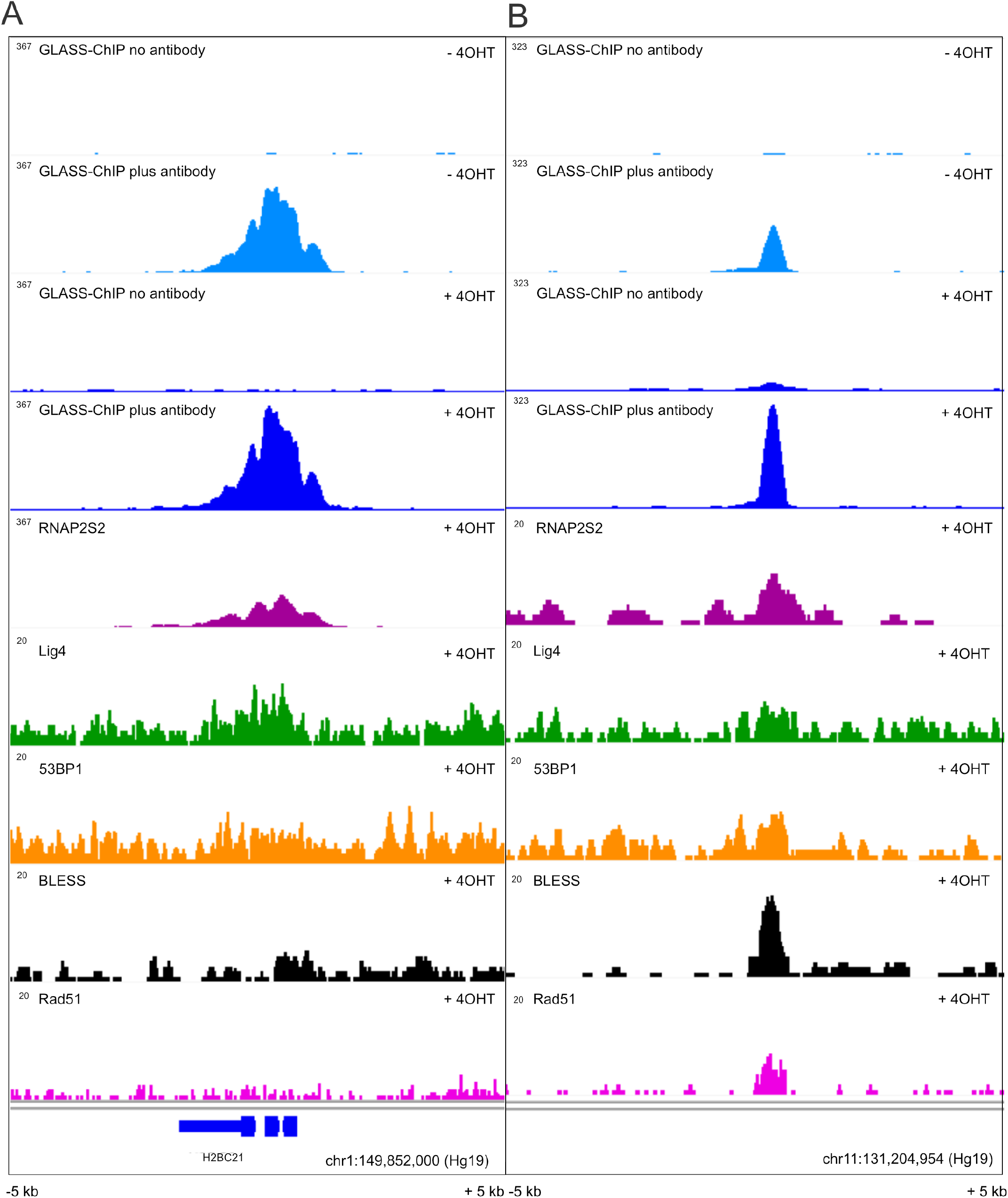
Examples of genome browser views of non-AsiSI sites where DNA-PK is bound in the human genome. (A,B) GLASS-ChIP (- and + 4-OHT), RNA Pol II phospho-S2 (RNAP2S2), Ligase 4, 53BP1, BLESS (DSBs), and Rad51 from this study as well as previous studies^1,2^.

**Figure S3.**
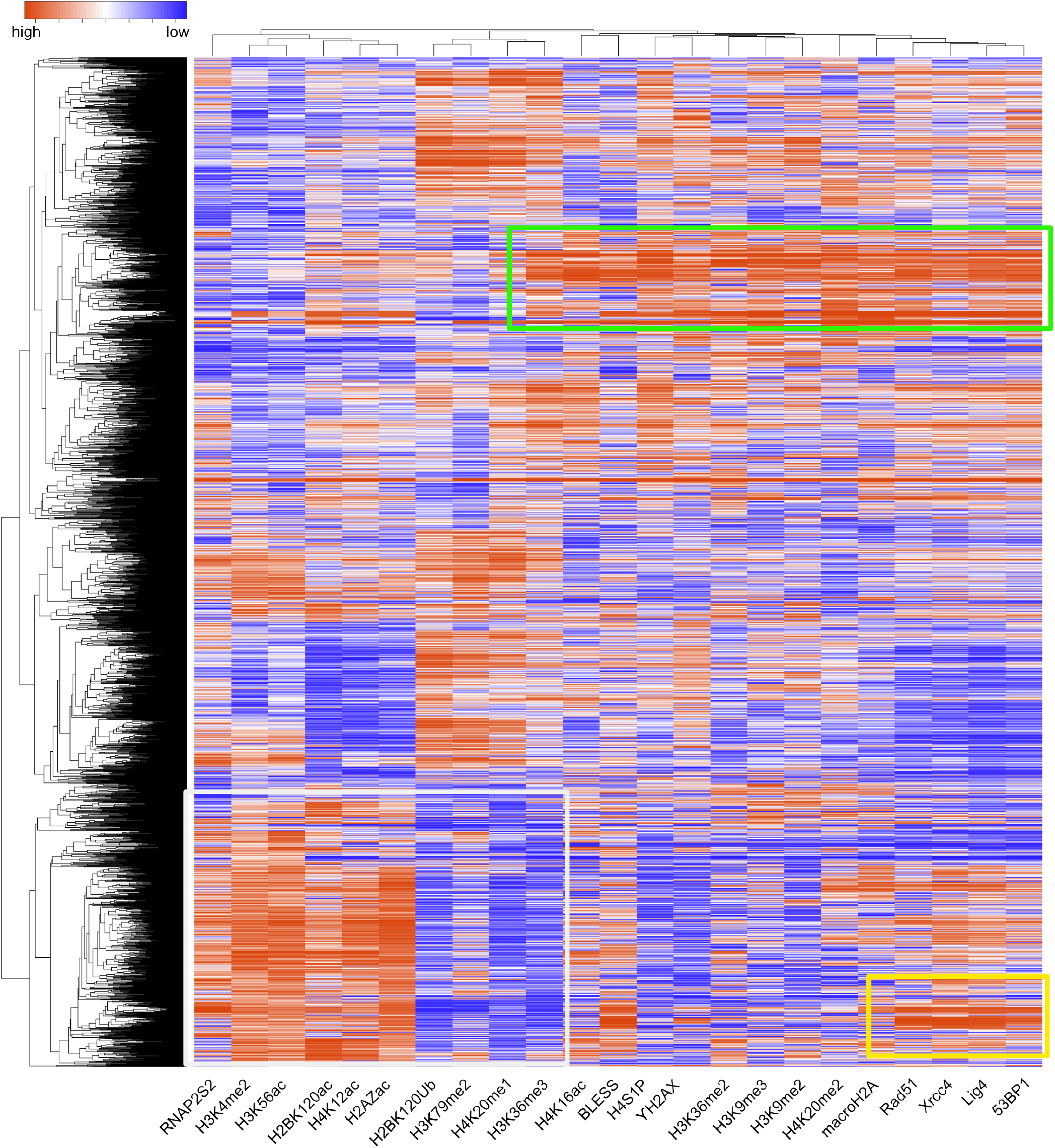
A subset of DNA-PKcs binding sites coincide with other DSB markers. The top 2500 binding sites from DNA-PKcs GLASS-ChIP were used to interrogate other DSB binding proteins and histone modifications from previous ChIP studies^1,2^. ChIP intensities for each factor/mark were used to generate rankings; these rankings were analyzed using unsupervised hierarchical clustering. Yellow box indicates one subset of sites where H4S1p, 53BP1, Rad51, Xrcc4, and Lig4 are enriched that contains many of the AsiSI sites; green box indicates other sites with DSB-associated marks also cosegregating with other chromatin marks; white indicates strongly anticorrelated marks present at a subset of GLASS-ChIP DNA-PKcs sites.

**Figure S4.**
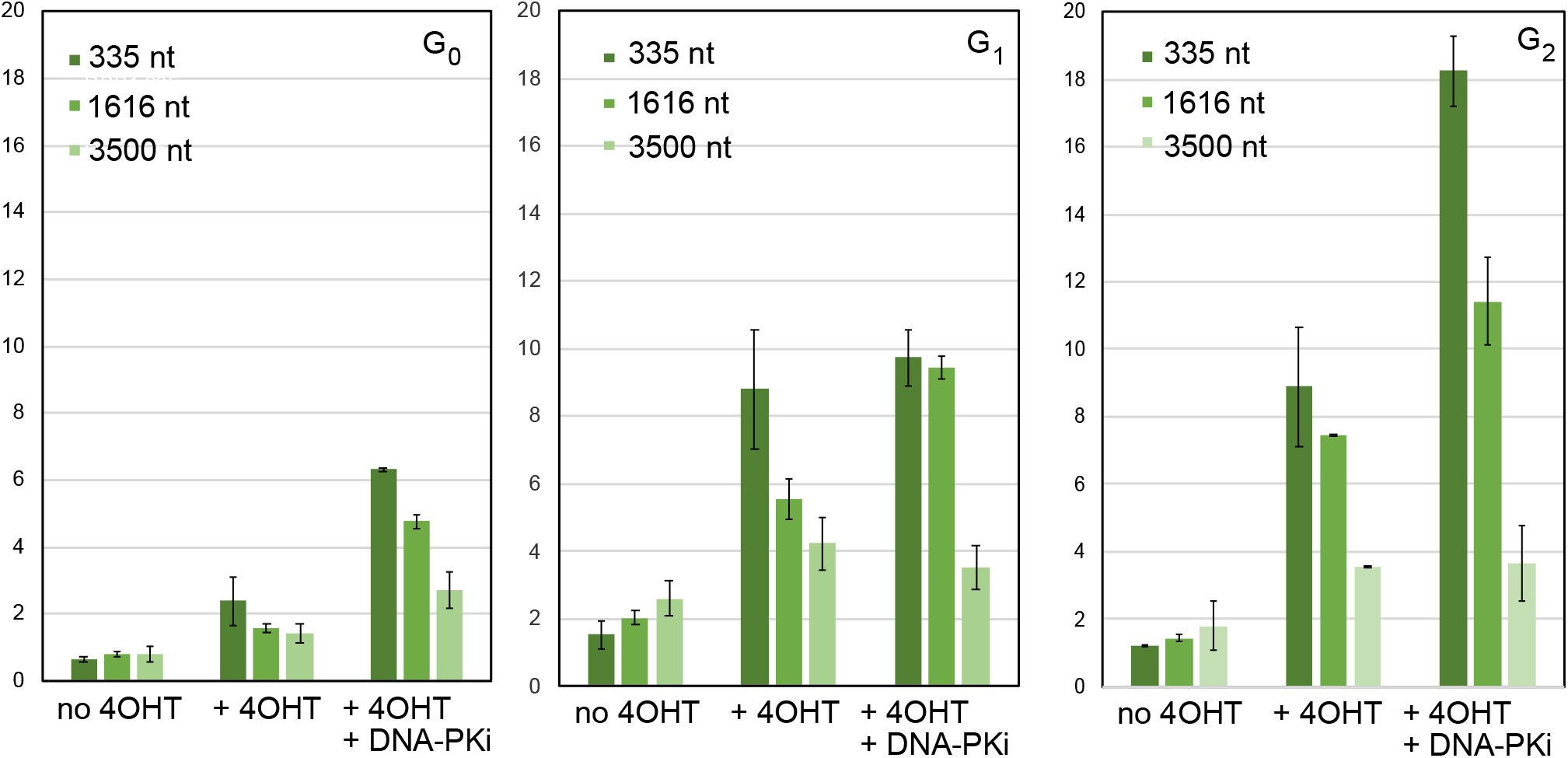
Resection at AsiSI breaks is observed in all cell cycle phases. Resection was monitored at various distances from the AsiSI site at the KYAT3 gene using a qPCR-based assay^3^ in U2OS cells in different cell cycle phases with 4-OHT and DNA-PKi as indicated.

**Figure S5.**
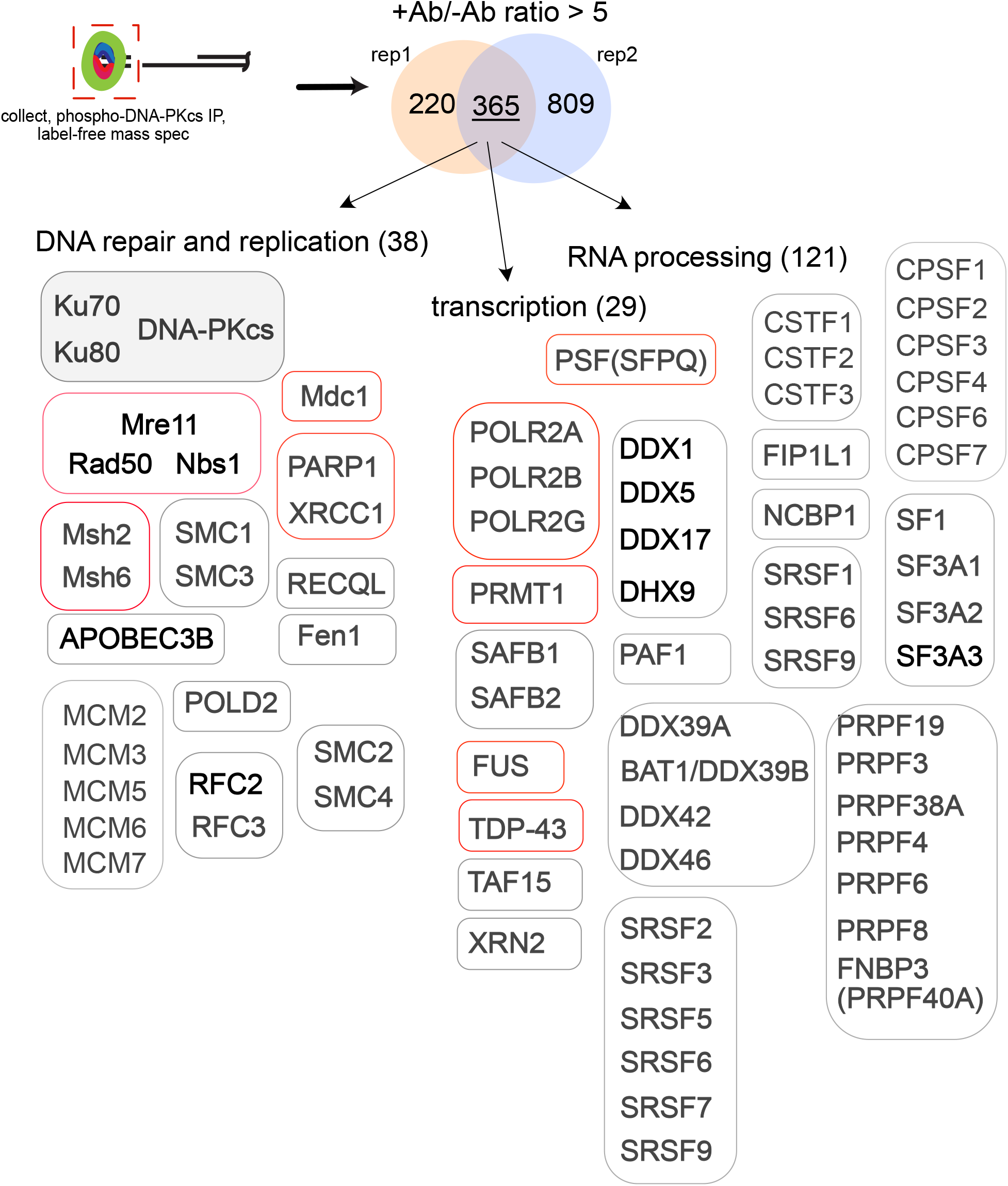
Mass Spectrometry identifies proteins associated with released DNA-PKcs GLASS-ChIP fragments. GLASS-ChIP was performed in two biological replicates, comparing -Ab to +Ab immunoprecipitations. Proteins were identified by label-free, quantitative mass spectrometry and targets with greater than 5-fold ratios between +Ab and -Ab were compared between the replicates as shown. Among the 365 targets are many DNA repair and replication factors as well as proteins involved in transcription and RNA processing. Factors previously known to associate with DNA-PK are highlighted in red. Complete list of factors in Table S2.

## References

1. Lees-Miller, J.P., Cobban, A., Katsonis, P., Bacolla, A., Tsutakawa, S.E., Hammel, M., Meek, K., Anderson, D.W., Lichtarge, O., Tainer, J.A., et al. (2020). Uncovering DNA-PKcs ancient phylogeny, unique sequence motifs and insights for human disease. Progress in Biophysics and Molecular Biology, S0079610720300973. 10.1016/j.pbiomolbio.2020.09.010.

2. Chang, H.H.Y., Pannunzio, N.R., Adachi, N., and Lieber, M.R. (2017). Non-homologous DNA end joining and alternative pathways to double-strand break repair. Nat Rev Mol Cell Biol 18, 495–506. 10.1038/nrm.2017.48.

3. Chen, C.-C., Feng, W., Lim, P.X., Kass, E.M., and Jasin, M. (2018). Homology-Directed Repair and the Role of BRCA1, BRCA2, and Related Proteins in Genome Integrity and Cancer. Annu Rev Cancer Biol 2, 313–336. 10.1146/annurev-cancerbio-030617-050502.

4. Symington, L.S. (2016). Mechanism and regulation of DNA end resection in eukaryotes. Crit Rev Biochem Mol Biol 51, 195–212. 10.3109/10409238.2016.1172552.

5. Paull, T.T. (2018). 20 Years of Mre11 Biology: No End in Sight. Mol. Cell 71, 419–427. 10.1016/j.molcel.2018.06.033.

6. Anand, R., Ranjha, L., Cannavo, E., and Cejka, P. (2016). Phosphorylated CtIP Functions as a Co-factor of the MRE11-RAD50-NBS1 Endonuclease in DNA End Resection. Mol Cell 64, 940–950. 10.1016/j.molcel.2016.10.017.

7. Anand, R., Jasrotia, A., Bundschuh, D., Howard, S.M., Ranjha, L., Stucki, M., and Cejka, P. (2019). NBS1 promotes the endonuclease activity of the MRE11-RAD50 complex by sensing CtIP phosphorylation. EMBO J. 38. 10.15252/embj.2018101005.

8. Deshpande, R.A., Lee, J.H., Arora, S., and Paull, T.T. (2016). Nbs1 Converts the Human Mre11/Rad50 Nuclease Complex into an Endo/Exonuclease Machine Specific for Protein-DNA Adducts. Mol Cell 64, 593–606. 10.1016/j.molcel.2016.10.010.

9. Deshpande, R.A., Myler, L.R., Soniat, M.M., Makharashvili, N., Lee, L., Lees-Miller, S.P., Finkelstein, I.J., and Paull, T.T. (2020). DNA-dependent protein kinase promotes DNA end processing by MRN and CtIP. Sci Adv 6, eaay0922. 10.1126/sciadv.aay0922.

10. Wang, W., Daley, J.M., Kwon, Y., Krasner, D.S., and Sung, P. (2017). Plasticity of the Mre11-Rad50-Xrs2-Sae2 nuclease ensemble in the processing of DNA-bound obstacles. Genes Dev 31, 2331–2336. 10.1101/gad.307900.117.

11. Reginato, G., Cannavo, E., and Cejka, P. (2017). Physiological protein blocks direct the Mre11-Rad50-Xrs2 and Sae2 nuclease complex to initiate DNA end resection. Genes Dev 31, 2325–2330. 10.1101/gad.308254.117.

12. Lam, I., and Keeney, S. (2014). Mechanism and regulation of meiotic recombination initiation. Cold Spring Harb Perspect Biol 7, a016634. 10.1101/cshperspect.a016634.

13. Brandsma, I., and Gent, D.C. (2012). Pathway choice in DNA double strand break repair: observations of a balancing act. Genome Integr 3, 9. 10.1186/2041-9414-3-9.

14. Kass, E.M., and Jasin, M. (Sep 10). Collaboration and competition between DNA double-strand break repair pathways. FEBS letters 584, 3703–3708. S0014-5793(10)00619-8 [pii] 10.1016/j.febslet.2010.07.057.

15. Iacovoni, J.S., Caron, P., Lassadi, I., Nicolas, E., Massip, L., Trouche, D., and Legube, G. (2010). High-resolution profiling of gammaH2AX around DNA double strand breaks in the mammalian genome. EMBO J 29, 1446–1457. 10.1038/emboj.2010.38.

16. Deshpande, R.A., and Paull, T.T. (2022). Characterization of DNA-PK-Bound End Fragments Using GLASS-ChIP. Methods Mol Biol 2444, 171–182. 10.1007/978-1-0716-2063-2_11.

17. Hoa, N.N., Shimizu, T., Zhou, Z.W., Wang, Z.Q., Deshpande, R.A., Paull, T.T., Akter, S., Tsuda, M., Furuta, R., Tsusui, K., et al. (2016). Mre11 Is Essential for the Removal of Lethal Topoisomerase 2 Covalent Cleavage Complexes. Mol Cell 64, 580–592. 10.1016/j.molcel.2016.10.011.

18. Shibata, A., Moiani, D., Arvai, A.S., Perry, J., Harding, S.M., Genois, M.-M., Maity, R., van Rossum-Fikkert, S., Kertokalio, A., Romoli, F., et al. (2014). DNA double-strand break repair pathway choice is directed by distinct MRE11 nuclease activities. Mol Cell 53, 7–18. 10.1016/j.molcel.2013.11.003.

19. Cohen, S., Puget, N., Lin, Y.-L., Clouaire, T., Aguirrebengoa, M., Rocher, V., Pasero, P., Canitrot, Y., and Legube, G. (2018). Senataxin resolves RNA:DNA hybrids forming at DNA double-strand breaks to prevent translocations. Nat Commun 9, 533. 10.1038/s41467-018-02894-w.

20. Aymard, F., Bugler, B., Schmidt, C.K., Guillou, E., Caron, P., Briois, S., Iacovoni, J.S., Daburon, V., Miller, K.M., Jackson, S.P., et al. (2014). Transcriptionally active chromatin recruits homologous recombination at DNA double-strand breaks. Nat. Struct. Mol. Biol. 21, 366–374. 10.1038/nsmb.2796.

21. Song, Z., Xie, Y., Guo, Z., Han, Y., Guan, H., Liu, X., Ma, T., and Zhou, P. (2019). Genome-wide identification of DNA-PKcs-associated RNAs by RIP-Seq. Sig Transduct Target Ther 4, 22. 10.1038/s41392-019-0057-6.

22. Bunch, H., Lawney, B.P., Lin, Y.-F., Asaithamby, A., Murshid, A., Wang, Y.E., Chen, B.P.C., and Calderwood, S.K. (2015). Transcriptional elongation requires DNA break-induced signalling. Nature Communications 6. 10.1038/ncomms10191.

23. Caron, P., Pankotai, T., Wiegant, W.W., Tollenaere, M.A.X., Furst, A., Bonhomme, C., Helfricht, A., de Groot, A., Pastink, A., Vertegaal, A.C.O., et al. (2019). WWP2 ubiquitylates RNA polymerase II for DNA-PK-dependent transcription arrest and repair at DNA breaks. Genes & Development 33, 684–704. 10.1101/gad.321943.118.

24. Clouaire, T., Rocher, V., Lashgari, A., Arnould, C., Aguirrebengoa, M., Biernacka, A., Skrzypczak, M., Aymard, F., Fongang, B., Dojer, N., et al. (2018). Comprehensive Mapping of Histone Modifications at DNA Double-Strand Breaks Deciphers Repair Pathway Chromatin Signatures. Molecular Cell 72, 250-262.e6. 10.1016/j.molcel.2018.08.020.

25. Zhou, Y., and Paull, T.T. (2015). Direct measurement of single-stranded DNA intermediates in mammalian cells by quantitative polymerase chain reaction. Analytical biochemistry 479, 48–50. 10.1016/j.ab.2015.03.025.

26. Shoaib, M., Chen, Q., Shi, X., Nair, N., Prasanna, C., Yang, R., Walter, D., Frederiksen, K.S., Einarsson, H., Svensson, J.P., et al. (2021). Histone H4 lysine 20 mono-methylation directly facilitates chromatin openness and promotes transcription of housekeeping genes. Nat Commun 12, 4800. 10.1038/s41467-021-25051-2.

27. Farooq, Z., Banday, S., Pandita, T.K., and Altaf, M. (2016). The many faces of histone H3K79 methylation. Mutat Res Rev Mutat Res 768, 46–52. 10.1016/j.mrrev.2016.03.005.

28. Sun, Z., Zhang, Y., Jia, J., Fang, Y., Tang, Y., Wu, H., and Fang, D. (2020). H3K36me3, message from chromatin to DNA damage repair. Cell Biosci 10, 9. 10.1186/s13578-020-0374-z.

29. Bar-Ziv, R., Voichek, Y., and Barkai, N. (2016). Chromatin dynamics during DNA replication. Genome Res. 26, 1245–1256. 10.1101/gr.201244.115.

30. Ma, Y., Kanakousaki, K., and Buttitta, L. (2015). How the cell cycle impacts chromatin architecture and influences cell fate. Front Genet 6, 19. 10.3389/fgene.2015.00019.

31. Armstrong, C., and Spencer, S.L. (2021). Replication-dependent histone biosynthesis is coupled to cell-cycle commitment. Proc. Natl. Acad. Sci. U.S.A. 118, e2100178118. 10.1073/pnas.2100178118.

32. tsai, R.Y.L., and Pederson, T. (2014). Connecting the nucleolus to the cell cycle and human disease. FASEB j. 28, 3290–3296. 10.1096/fj.14-254680.

33. Zhou, Y., and Paull, T.T. (2013). DNA-dependent Protein Kinase Regulates DNA End Resection in Concert with Mre11-Rad50-Nbs1 (MRN) and Ataxia Telangiectasia-mutated (ATM). J Biol Chem 288, 37112–37125. 10.1074/jbc.M113.514398.

34. Littlewood, T.D., Hancock, D.C., Danielian, P.S., Parker, M.G., and Evan, G.I. (1995). A modified oestrogen receptor ligand-binding domain as an improved switch for the regulation of heterologous proteins. Nucleic Acids Res 23, 1686–1690. 10.1093/nar/23.10.1686.

35. Liu, Y., Zou, R.S., He, S., Nihongaki, Y., Li, X., Razavi, S., Wu, B., and Ha, T. (2020). Very fast CRISPR on demand. Science 368, 1265–1269. 10.1126/science.aay8204.

36. Zou, R.S., Marin-Gonzalez, A., Liu, Y., Liu, H.B., Shen, L., Dveirin, R.K., Luo, J.X.J., Kalhor, R., and Ha, T. (2022). Massively parallel genomic perturbations with multi-target CRISPR interrogates Cas9 activity and DNA repair at endogenous sites. Nat Cell Biol 24, 1433–1444. 10.1038/s41556-022-00975-z.

37. Cannavo, E., and Cejka, P. (2014). Sae2 promotes dsDNA endonuclease activity within Mre11-Rad50-Xrs2 to resect DNA breaks. Nature 514, 122–125. 10.1038/nature13771.

38. Kim, J.-S., Krasieva, T.B., Kurumizaka, H., Chen, D.J., Taylor, A.M.R., and Yokomori, K. (2005). Independent and sequential recruitment of NHEJ and HR factors to DNA damage sites in mammalian cells. The Journal of Cell Biology 170, 341–347. 10.1083/jcb.200411083.

39. Yang, G., Liu, C., Chen, S.-H., Kassab, M.A., Hoff, J.D., Walter, N.G., and Yu, X. (2018). Super-resolution imaging identifies PARP1 and the Ku complex acting as DNA double-strand break sensors. Nucleic Acids Res. 46, 3446–3457. 10.1093/nar/gky088.

40. Kochan, J.A., Desclos, E.C.B., Bosch, R., Meister, L., Vriend, L.E.M., van Attikum, H., and Krawczyk, P.M. (2017). Meta-analysis of DNA double-strand break response kinetics. Nucleic Acids Research 45, 12625–12637. 10.1093/nar/gkx1128.

41. Jette, N., and Lees-Miller, S.P. (2015). The DNA-dependent protein kinase: A multifunctional protein kinase with roles in DNA double strand break repair and mitosis. Progress in biophysics and molecular biology 117, 194–205. 10.1016/j.pbiomolbio.2014.12.003.

42. Reddy, Y.V., Ding, Q., Lees-Miller, S.P., Meek, K., and Ramsden, D.A. (2004). Non-homologous end joining requires that the DNA-PK complex undergo an autophosphorylation-dependent rearrangement at DNA ends. J Biol Chem 279, 39408–39413. 10.1074/jbc.M406432200M406432200[pii].

43. Ding, Q., Reddy, Y.V., Wang, W., Woods, T., Douglas, P., Ramsden, D.A., Lees-Miller, S.P., and Meek, K. (2003). Autophosphorylation of the catalytic subunit of the DNA-dependent protein kinase is required for efficient end processing during DNA double-strand break repair. Mol Cell Biol 23, 5836–5848.

44. Block, W.D., Yu, Y., Merkle, D., Gifford, J.L., Ding, Q., Meek, K., and Lees-Miller, S.P. (2004). Autophosphorylation-dependent remodeling of the DNA-dependent protein kinase catalytic subunit regulates ligation of DNA ends. Nucleic acids research 32, 4351–4357. 10.1093/nar/gkh76132/14/4351[pii].

45. Lee, K.J., Saha, J., Sun, J., Fattah, K.R., Wang, S.C., Jakob, B., Chi, L., Wang, S.Y., Taucher-Scholz, G., Davis, A.J., et al. (2016). Phosphorylation of Ku dictates DNA double-strand break (DSB) repair pathway choice in S phase. Nucleic acids research 44, 1732–1745. 10.1093/nar/gkv1499.

46. Calsou, P., Frit, P., Humbert, O., Muller, C., Chen, D.J., and Salles, B. (1999). The DNA-dependent protein kinase catalytic activity regulates DNA end processing by means of Ku entry into DNA. J Biol Chem 274, 7848–7856.

47. de Vries, E., van Driel, W., Bergsma, W.G., Arnberg, A.C., and van der Vliet, P.C. (1989). HeLa nuclear protein recognizing DNA termini and translocating on DNA forming a regular DNA-multimeric protein complex. J Mol Biol 208, 65–78. 10.1016/0022-2836(89)90088-0.

48. Paillard, S., and Strauss, F. (1991). Analysis of the mechanism of interaction of simian Ku protein with DNA. Nucleic Acids Res 19, 5619–5624. 10.1093/nar/19.20.5619.

49. Woodard, R.L., Anderson, M.G., and Dynan, W.S. (1999). Nuclear Extracts Lacking DNA-dependent Protein Kinase Are Deficient in Multiple Round Transcription. J. Biol. Chem. 274, 478–485. 10.1074/jbc.274.1.478.

50. Nock, A., Ascano, J.M., Jones, T., Barrero, M.J., Sugiyama, N., Tomita, M., Ishihama, Y., and Malik, S. (2009). Identification of DNA-dependent Protein Kinase as a Cofactor for the Forkhead Transcription Factor FoxA2. Journal of Biological Chemistry 284, 19915–19926. 10.1074/jbc.M109.016295.

51. Salifou, K., Burnard, C., Basavarajaiah, P., Grasso, G., Helsmoortel, M., Mac, V., Depierre, D., Franckhauser, C., Beyne, E., Contreras, X., et al. (2021). Chromatin-associated MRN complex protects highly transcribing genes from genomic instability. Sci Adv 7. 10.1126/sciadv.abb2947.

52. Ingram, S.P., Warmenhoven, J.W., Henthorn, N.T., Smith, E.A.K., Chadwick, A.L., Burnet, N.G., Mackay, R.I., Kirkby, N.F., Kirkby, K.J., and Merchant, M.J. (2019). Mechanistic modelling supports entwined rather than exclusively competitive DNA double-strand break repair pathway. Sci Rep 9, 6359. 10.1038/s41598-019-42901-8.

53. Chen, S., Zhou, Y., Chen, Y., and Gu, J. (2018). fastp : an ultra-fast all-in-one FASTQ preprocessor (Bioinformatics) 10.1101/274100.

54. Li, H., and Durbin, R. (2010). Fast and accurate long-read alignment with Burrows–Wheeler transform. Bioinformatics 26, 589–595. 10.1093/bioinformatics/btp698.

55. Li, H., Handsaker, B., Wysoker, A., Fennell, T., Ruan, J., Homer, N., Marth, G., Abecasis, G., Durbin, R., and 1000 Genome Project Data Processing Subgroup (2009). The Sequence Alignment/Map format and SAMtools. Bioinformatics 25, 2078–2079. 10.1093/bioinformatics/btp352.

56. Zhang, Y., Liu, T., Meyer, C.A., Eeckhoute, J., Johnson, D.S., Bernstein, B.E., Nusbaum, C., Myers, R.M., Brown, M., Li, W., et al. (2008). Model-based analysis of ChIP-Seq (MACS). Genome Biol. 9, R137. 10.1186/gb-2008-9-9-r137.

57. Babicki, S., Arndt, D., Marcu, A., Liang, Y., Grant, J.R., Maciejewski, A., and Wishart, D.S. (2016). Heatmapper: web-enabled heat mapping for all. Nucleic Acids Res 44, W147–153. 10.1093/nar/gkw419.

58. Ryu, S.W., Stewart, R., Pectol, C., Ender, N., Wimalarathne, O., Lee, J.-H., Zanini, C.P., Harvey, A., Huibregtse, J., Mueller, P., et al. (2019). Comprehensive identification of HSP70/HSC70 Chaperone Clients in Human Cells (Molecular Biology) 10.1101/865030.

59. Lee, J.-H., Mand, M.R., Kao, C.-H., Zhou, Y., Ryu, S.W., Richards, A.L., Coon, J.J., and Paull, T.T. (2018). ATM directs DNA damage responses and proteostasis via genetically separable pathways. Sci Signal 11. 10.1126/scisignal.aan5598.

## References cited

1. Aymard, F. et al. Transcriptionally active chromatin recruits homologous recombination at DNA double-strand breaks. Nat. Struct. Mol. Biol. 21, 366–374 (2014)

2. Clouaire, T. et al. Comprehensive Mapping of Histone Modifications at DNA Double-Strand Breaks Deciphers Repair Pathway Chromatin Signatures. Molecular Cell 72, 250–262.e6 (2018).

3. Zhou, Y., Caron, P., Legube, G. & Paull, T. T. Quantitation of DNA double-strand break resection intermediates in human cells. Nucleic acids research 42, e19 (2014).

